# Thalamus drives two complementary input strata of the neocortex in parallel

**DOI:** 10.1101/524298

**Authors:** R. Egger, R.T. Narayanan, D. Udvary, A. Bast, J.M. Guest, S. Das, C.P.J. de Kock, M. Oberlaender

## Abstract

Sensory information enters the neocortex via thalamocortical axons that define the major ‘input’ layer 4. The same thalamocortical axons, however, additionally innervate the deep ‘output’ layers 5/6. How such bistratification impacts cortical processing remains unknown. Here, we find a class of neurons that cluster specifically around thalamocortical axons at the layer 5/6 border. We show that these border *stratum cells* are characterized by extensive horizontal axons, that they receive strong convergent input from the thalamus, and that this input is sufficient to drive reliable sensory-evoked responses, which precede those in layer 4. These cells are hence strategically placed to amplify and relay thalamocortical inputs across the cortical area, for example to drive the fast onsets of cortical output patterns. Layer 4 is therefore not the sole starting point of cortical processing. Instead, parallel activation of layer 4 and the border stratum is necessary to broadcast information out of the neocortex.

## Introduction

The mammalian neocortex is required for higher-order brain functions, such as sensory perception and cognition, and hence for the transformation of information from the environment into behavior. Such information, for example as evoked by photo- or mechanoreceptor cells at the periphery of the visual or somatosensory system, enters the neocortex in form of a synchronous volley of excitation, which is provided by relay cells that are located in sensory system-specific primary nuclei of the thalamus ^1^. Despite several species- and sensory system-specific differences, thalamocortical axons of relay cells terminate in general most densely in layer 4 of the respective primary sensory areas of the neocortex ^2^. For decades, concepts about the neocortex thus focused on layer 4 as its main input site, and starting point of sensory information processing ^3^. However, anatomical studies in macaques ^4^, cats ^5^, and rodents ^6^ indicate that the very same thalamocortical axons give rise to a second innervation peak at specific depth locations in the deeper layers 5 and/or 6. These layers represent the main output site of the neocortex, as they comprise long-range projection neurons that innervate subcortical brain structures ^2^.

The relevance of bistratified thalamocortical input for cortical information processing – and in particular of the deep input stratum within the output layers – remains poorly understood ^7^. However, this knowledge is fundamental for deducing the logic of intracortical signal flow, for revealing the origin of cell type- and layer-specific neuronal activity patterns, and for constraining hypotheses of cortical circuit organization. So far, to our knowledge, no study has yet systematically investigated the principles by which a deep thalamocortical input stratum contributes to the overall sensory-evoked cortical excitation. It remains therefore unknown whether and how different types of neurons in the output layers can be driven directly by thalamocortical input, and if that would be the case, how responses in the deep input stratum affect those in the upper layers and vice versa.

Here, we address these questions in the whisker somatosensory system of the rat ^8^. Tactile information from whisker touch enters the neocortex via relay cells from the ventral posterior medial nucleus (VPM) of the thalamus ^2^. Axons from these relay cells delineate layer 4 of the whisker-related part of the primary somatosensory cortex (wS1). Within layer 4, excitatory spiny neurons cluster around the dense terminal fields of the VPM axons. Upon sensory stimulation, this major thalamorecipient population gives rise to recurrent excitation within layer 4 and feed-forward excitation to the superficial layers 2/3 – a canonical organizational principle of all sensory cortices ^3^. VPM axons show a second, less dense innervation peak in the deeper layers ^6^. The VPM-to-wS1 pathway in rodents hence represents an ideal model system to elucidate the relevance of bistratified thalamocortical input for cortical sensory information processing.

Combining *in vivo* recordings with morphological reconstructions, optogenetic input mappings, pharmacological manipulations, and simulations of cortical signal flow, we reveal that – similar to the organization of layer 4 – thalamocortical inputs converge strongly onto a population of corticocortical cells that is strategically placed around the terminal fields of VPM axons at the layer 5/6 border. Upon sensory stimulation, this deep thalamorecipient pathway is activated first, evoking a stream of excitation that spreads horizontally across the deep layers, and which thereby precedes the vertical stream of signal flow through the canonical layer 4 to layers 2/3 pathway. Neuronal responses in primary sensory cortices may thus be regarded as a superposition of inputs from two simultaneously active primary thalamocortical pathways, which likely complement each other to ensure that intracortical computations are reliably transformed into cortical output patterns.

## Results

### Corticocortical cells cluster around deep layer thalamocortical axons

We compared the soma, dendrite, and axon distributions of all major excitatory cortical cell types ^2^ with the distributions of thalamocortical axons, and precise measurements of the cytoarchitectonic layer borders **(Fig. S1)** – anatomical data that we had systematically collected for one set of experimental conditions over the past decade in rat VPM and wS1 ^6, 9, 10, 11, 12, 13, 14^. The comparison revealed that the peak density of VPM axons in the deep layers coincides with the cytoarchitectonic border between layers 5 and 6 **(Fig. 1A)**. Moreover, the soma depth distribution of layer 6 corticocortical (i.e., intratelencephalic) neurons matches the vertical extent of the deep layer VPM axon density peak **(Fig. 1B)**. Neurons of this class are hence not restricted to layer 6, but equally abundant in lower layer 5 and upper layer 6.

**Figure 1:**
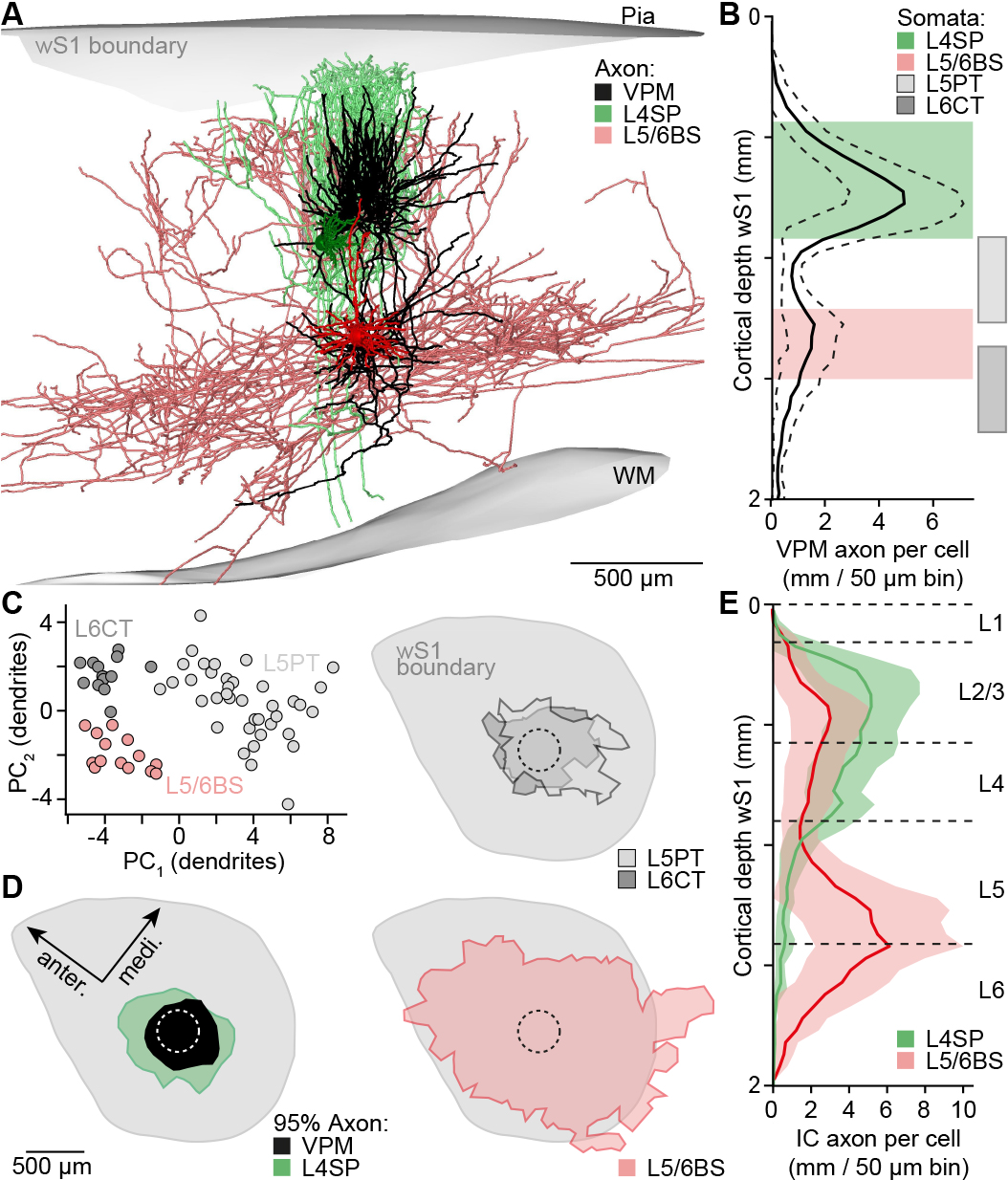
Structural basis of two thalamocortical input strata. **A)** Examples of *in vivo* labeled neurons in the whisker-related part of rat primary somatosensory cortex (wS1): layer 4 spiny neuron (L4SP), corticocortical neuron at the L5/6 border stratum (L5/6BS), and the intracortical (IC) part of the axon from a relay cell in the ventral posterior medial nucleus of the thalamus (VPM). **B)** Soma, dendrite and axon distributions of induvial neurons (n=191) were compared with 50 μm precision ^9^. Somata of L4SP (n=37) and L5/6BS cells (n=14) cluster around the two innervation peaks of VPM axons (n=14, mean ± STD). Somata of L5/6BS cells intermingle with those of the subcortically projecting pyramidal tract (L5PT, n=38) and corticothalamic (L6CT, n=13) neurons. **C)** Principal components (PC_1/2_) of dendritic features that discriminate between excitatory cell types in the deep layers ^12, 13^ (representing the cells in panel B). **D)** Horizontal axon extent (95% iso-contours) of VPM (n=14), L4SP (n=14), L5PT (n=7), L6CT (n=11) and L5/6BS (n=9) neurons. Top views onto wS1. **E)** Vertical distributions of L4SP and L5/6BS axons (same cells as in panel D) vs. cytoarchitectonic layer borders ^11^.

These layer 5/6 corticocortical neurons can be easily distinguished from the subcortically projecting cortical output neurons that are found at the same depth: layer 5 pyramidal tract and layer 6 corticothalamic neurons. First, by the characteristic morphology of their apical dendrites **(Fig. 1C)**, which terminate in layer 4 without forming a tuft. This property also distinguishes them from polymorphic corticocortical neurons in deeper regions of layer 6 ^12^. Second, by their extensive horizontally projecting axons **(Fig. 1D)**, which can span across the deep layers of the entire cortical area **(Fig. 1E)**. Our data reveal that analogously to the organization of excitatory populations in the major thalamorecipient layer, neurons with dendrite morphologies similar to those in layer 4, but with complementary intracortical axon projection patterns, cluster around thalamocortical axons in the deep layers. Together, VPM axons and layer 5/6 corticocortical neurons thus provide the structural basis of a deep thalamorecipient pathway for sensory-evoked signal flow, which we subsequently refer to as the ‘*layer 5/6 border stratum*’ ^4^.

### Thalamocortical inputs converge strongly onto border stratum cells

Functional and anatomical studies in the rodent somatosensory ^7, 15^ and visual systems ^16^ suggest that neurons of all excitatory cell types that are located at the depth of the border stratum can receive direct input from the respective primary thalamic nuclei. We therefore quantified the degree to which border stratum cells (i.e., layer 5/6 corticocortical) in rat wS1 form monosynaptic connections with VPM axons, and investigated whether these synapses are functional under *in vivo* conditions. We expressed light-gated ion channels and a fluorescent marker within the thalamocortical synapses by injecting an adeno-associated virus into the VPM **(Fig. 2A)**. Light-evoked and sensory-evoked action potential (AP) responses of individual wS1 neurons were obtained via cell-attached recordings in anesthetized rats **(Fig. 2B)**. Following the recordings, neurons were filled *in vivo* with biocytin, which allowed for post-hoc reconstruction and classification of the neurons’ morphology **(Fig. 2C)**, and detection of the putative thalamocortical synapses along the dendrites of the recorded neurons **(Fig. 2D)**.

**Figure 2:**
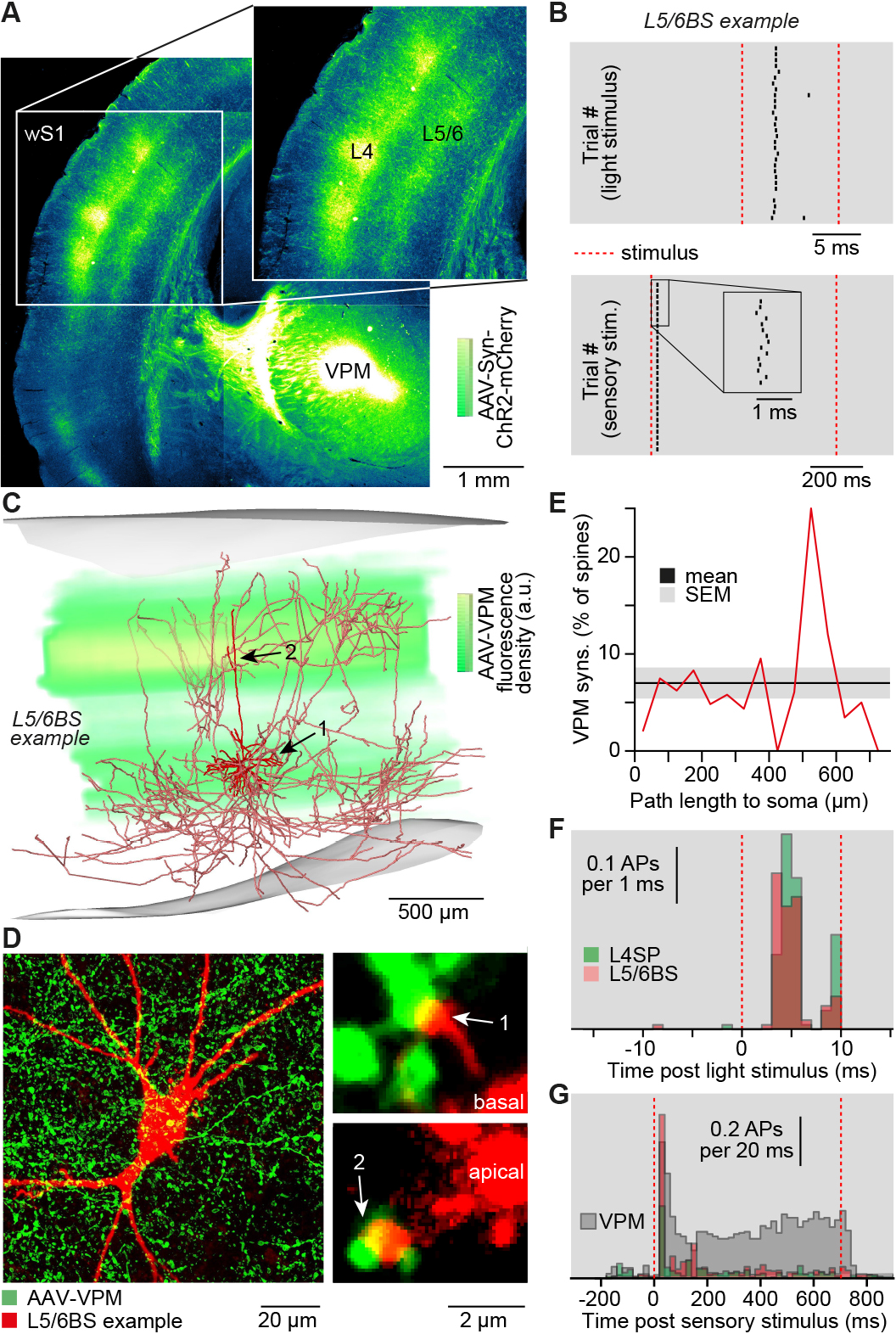
Synaptic basis of two thalamocortical input strata. **A)** Coronal brain section (50 μm) after injection of an adeno-associated virus (AAV) into the VPM, which expresses channel rhodopsin (ChR2) and a fluorescent marker (mCherry) in thalamocortical (TC) synapses. **B)** Example of cell-attached *in vivo* recording in wS1 of AAV-injected brain. Ticks represent APs in response to a 10 ms flash of green light onto the cortical surface (upper panel), and a 700 ms airpuff onto the whiskers (lower panel). **C)** Reconstruction of the L5/6BS cell shown in panel B, superimposed with quantification of AAV labeling. **D)** Confocal images of the L5/6BS cell shown in panel C. Putative TC synapses were identified as contacts between VPM boutons and dendritic spines. **E)** Fraction of spines (n=4789) along the dendrites of the L5/6BS cell shown in panel B-D that are contacted by VPM boutons. **F)** Post-stimulus-time-histograms (PSTHs) of light-evoked APs in L4SP and L5/6BS cells (mean ± STD of AP onset: 4.6±0.7 ms, n=4 vs. 4.4±0.8 ms, n=4). **G)** PSTHs of airpuff-evoked APs in L4SP, L5/6BS (same cells as in panel F) and VPM cells (n=7).

The experiments revealed that 7±2% of the spines along the dendrites of a border stratum cell receive input from the VPM **(Fig. 2E)**. Similar fractions (10±4%) and dendritic distributions of VPM synapses have been reported previously for spiny neurons in layer 4 of rat wS1 ^17^. Thus, the excitatory populations that define the respective postsynaptic parts of the two thalamocortical input strata – layer 4 spiny neurons and layer 5/6 border stratum cells – receive similar relative amounts of VPM input. Supporting these anatomical observations, light stimulation of VPM synapses elicited APs in the morphologically identified border stratum cells. The responses were equally reliable and as fast as those in layer 4 spiny neurons **(Fig. 2F)**. The same border stratum cells also responded to sensory stimuli, as evoked by a low pressure airpuff that deflects all whiskers caudally. Under these conditions, sensory responses in border stratum cells were more reliable compared to those of spiny neurons in layer 4, and even rivaled the reliability of relay cells in the VPM **(Fig. 2G)**. Our data reveal that analogously to layer 4 **(Fig. S2)**, the strategic location of border stratum cells results in strong convergent input from primary thalamocortical axons, which provides the synaptic basis of a deep thalamorecipient pathway for sensory-evoked signal flow.

### Border stratum cells respond first to sensory stimuli

Whole-cell recordings in rodent wS1 ^18^ and V1 ^16^ indicate that deep layer corticocortical neurons have intrinsic physiological properties that render them as highly excitable when compared to corticothalamic neurons that are found at the same depth. Together with our observation of strong convergence of thalamocortical axons, this suggests that synaptic input from these fibers may be sufficient to drive reliable sensory-evoked responses in border stratum cells (e.g. those shown in **Fig. 2G**). This hypothesis is supported by several studies which showed that response onsets (i.e., latency to first AP) of deep layer neurons can rival, and even precede those in layer 4 ^7, 10, 19^. To quantitatively test this hypothesis, we measured the additional path length between the border stratum and layer 4 that APs need to travel along VPM axons. Combined with conduction velocity measurements ^20^, the analysis predicted that sensory-evoked excitation reaches the border stratum 2 to 5 ms (3.0±1.7 ms) earlier than layer 4 **(Fig. 3A)**.

**Figure 3:**
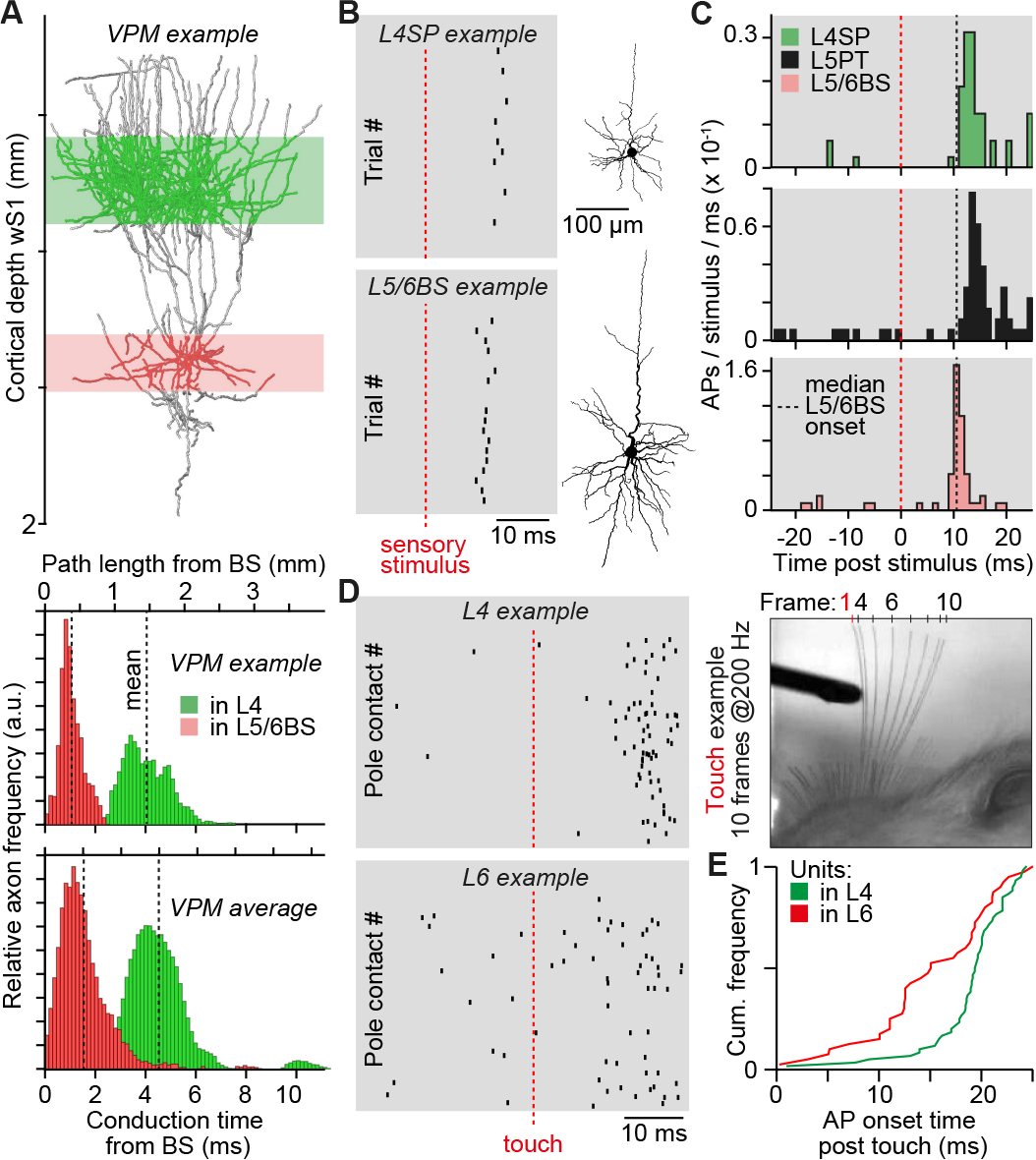
Functional basis of two thalamocortical input strata. **A)** Example of *in vivo* labeled VPM axon (upper panel), whose path length distribution was quantified with respect to the deepest location of the border stratum (L5/6BS), and multiplied with the IC conduction velocity (0.33 m/s ^20^) of TC axons (center panel). Average conduction time of VPM axons (n=14) to the border stratum and layer 4 (lower panel). **B)** AP responses evoked by principal whisker (PW) deflections in exemplary L4SP and L5/6BS cells. **C)** PSTHs of PW-evoked APs in morphologically identified L4SP (n=8), L5PT (n=9) and L5/6BS (n=6) cells. **D)** Example of simultaneously recorded single units in layer 4 and upper layer 6 (~1.6 mm recording depth), which show reliable AP responses after PW contact with a pole during exploratory whisking (right panel: whisker positions after exemplary touch). **E)** Distribution of touch-evoked AP onsets across animals (n=3) in layer 4 and upper layer 6.

To test the prediction, we recorded and labeled excitatory neurons across all layers of wS1 in anesthetized rats. These experiments allowed to precisely control the stimulus onset by deflecting individual whiskers with a piezoelectric bimorph ^10^, and to recover the morphological cell type of the recorded neurons **(Fig. 3B)**. We found that responses in the deep layers to deflections of the whisker that was somatotopically aligned with the recording site – the principal whisker – were largely restricted to the populations of border stratum and pyramidal tract neurons **(Fig. S3)**. Similar to the multi-whisker stimulations by airpuff, single whisker deflections evoked AP responses that were more reliable in border stratum cells when compared to layer 4 neurons **(Fig. 3C)**. Response onsets of border stratum cells (median/25^th^/75^th^ percentile: 11.2/10.3/12.4 ms) preceded those in all other excitatory cell types – including layer 4 – matching the path length-based delay predictions (14.3/13.3/18.4 ms; two-sided Mann-Whitney U-test: difference: −3.3, 95% CI [−4.0, −2.6], U = 1096, p < 10^−10^).

We further tested the delay predictions by simultaneously recording AP responses in layer 4 and upper layer 6 of head-fixed, behaving rats. We implanted linear silicon probes with equally-spaced electrodes that spanned across the depth of wS1. This allowed to record the AP activity of several single units during awake conditions, and to determine the units’ respective depth locations with ±50 μm precision. When animals explored their environment by rhythmically moving the principal whisker – all other whiskers were trimmed – sensory input was provided by whisker contact with a pole that was placed within range. AP responses in upper layer 6 preceded those in layer 4 of the same animal **(Fig. 3D)**. Across animals, the average AP onset in layer 4 was hence significantly delayed compared to layer 6 (Kolmogorov-Smirnov two-sample test: D_60,40_ = 0.425, p < 0.01), on average by 4 ms **(Fig. 3E)**. Our data reveal that inputs from primary thalamic axons can reliably drive fast APs in border stratum cells, providing the functional basis of a deep thalamorecipient pathway for sensory-evoked signal flow.

### Manipulating border stratum cells affects broad tuning of cortical output

For the present conditions of single whisker deflections in anesthetized rats, AP responses of pyramidal tract neurons occurred near simultaneous with those in layer 4, and hence consistently later than in border stratum cells – approximately 3-4 ms (14.3/13.6/16.2 ms). One of the functions of the border stratum pathway could thus be that it is involved in driving cortical output patterns, whose onsets thereby rival those in layer 4. In further supported of this hypothesis is the characteristic property of pyramidal tract neurons to respond to a broader range of stimuli compared to their thalamocortical input neurons ^21^. In case of wS1, pyramidal tract neurons can respond similarly fast to stimulations of several whiskers ^22^, even if their dendrites are located hundreds of micrometers away from the terminal fields of those VPM axons that provide the respective thalamocortical input ^23^. The extensive horizontally projecting axons in the deep layers, in combination with the earliest and reliable AP responses, hence render border stratum cells as ideal candidates that could contribute to the fast onsets and broadly tuned characteristics of cortical output patterns.

To test this hypothesis, we combined pharmacological injections of the GABAA agonist muscimol with cell-attached recordings in anesthetized rats **(Fig. 4A)**. Injection pipettes were positioned at the layer 5/6 border of wS1 by quantifying local field potentials (LFPs) at different cortical depths **(Fig. S4)**. The LFP recordings allowed mapping of the principal whisker that corresponded to the location of the injection site ^24^ – here referred to as the ‘manipulated whisker’. Cell-attached recordings were performed in layer 5, approximately 1 millimeter away from the injection site. This distance assured that axons from border stratum cells, but not from other cell types that are affected by the pharmacology, overlap with the recording site **(Fig. 4B)**. Pyramidal tract neurons were identified as those that responded to deflections of several individual whiskers ^22^ – including the manipulated whisker **(Fig. 4C)**. After muscimol injections, fast responses evoked by the manipulated whisker were abolished in any of the recorded neurons **(Fig. 4D)**. In contrast, whiskers that were not somatotopically aligned with the injection site (e.g. the principal whisker at the recording site) maintained their ability to evoke reliable and fast AP responses **(Fig. 4E)**.

**Figure 4:**
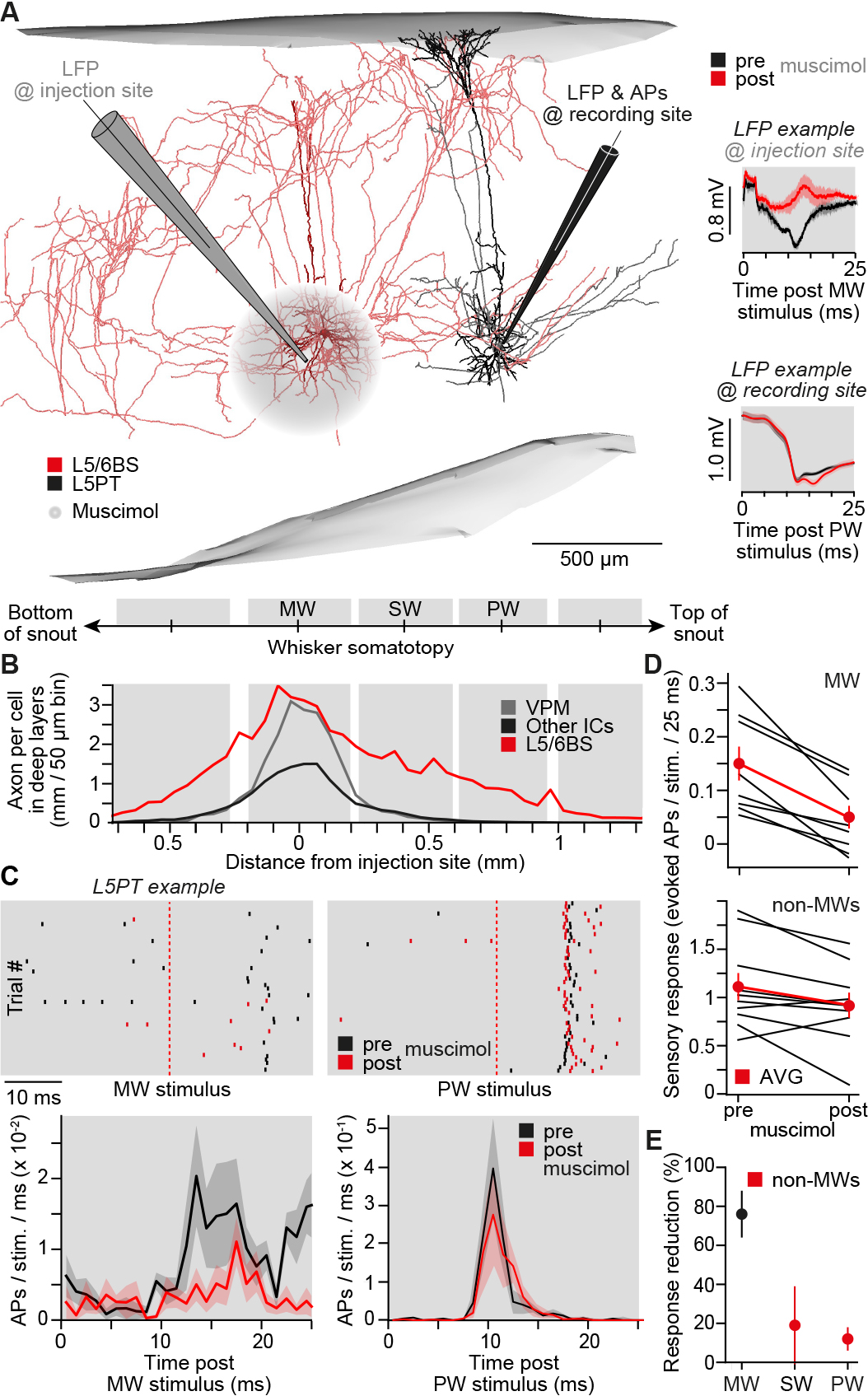
Onsets of broadly tuned cortical output patterns are driven by L5/6BS cells. **A)** The somatotopy of rat wS1 ^9^ in combination with whisker-evoked local field potential (LFP) measurements ^24^, allowed placing of muscimol injection and recording pipettes such that the respective PWs were separated by one whisker. Here the PW at the recording site is B2, the manipulated whisker (MW) is D2, and the separating whisker (SW) is C2. Left panel: L5/6BS and L5PT cells labeled in the same animal illustrate pharmacology experiments. Right panels: example LFPs before and after muscimol injections. **B)** Axonal extent in the deep layers from neurons located in the barrel column that represents the MW. **C)** Exemplary AP responses evoked by MW and PW deflections and PSTHs across cells (mean ± SEM; MW: n=8, PW: n=5). **D)** Response per cell to deflections of the MW (n=8, Wilcoxon rank-sum test: median = 0.095, 95% CI [0.05, 0.16], W=36, p=0.008) and non-MWs (PW & SW, n=5 & 5, Wilcoxon rank-sum test: median = 0.18, 95% CI [6×10^−5^, 0.38], W=47.5, p=0.05) before and after muscimol injections. Mean ± SEM. **E)** Effect of muscimol on L5PT responses to deflections of the MW and non-MWs.

### Border stratum cells provide an on-switch for cortical output

The pharmacological manipulations suggest that border stratum cells are necessary to drive the fast component of broadly tuned responses in pyramidal tract neurons. However, the high degree of recurrence in cortical networks, as well as non-linear mechanisms of synaptic and/or dendritic integration, pose a general challenge to infer causality between manipulations and the resultant alterations of neuronal AP responses. To address these issues, we developed a model that allows performing simulations that mimic the specific conditions of our *in vivo* pharmacology experiments at synaptic, cellular, and network levels, in the following referred to as *in silico* experiments. A link to download the model and simulation routines, and a detailed description and validation of all parameters, is provided in the **SI**. In brief, we embedded the morphology of an *in vivo* labeled pyramidal tract neuron into a previously reported anatomically realistic network model of rat wS1 ^25^. The embedding provided structural constraints about which neurons, depending on their respective cell type and location within VPM and wS1 (i.e., neurons represent *in vivo* labeled morphologies), can in principle form synaptic connections with the pyramidal tract neuron **(Fig. 5A)**, and where along its dendrites **(Fig. 5B)**. Combining the network model with cell type-specific AP measurements – acquired during conditions that were consistent with those of our pharmacology experiments ^10, 23, 26^ – provided functional constraints about which of the structurally possible connections can in principle provide input to the pyramidal tract neuron, depending on the identity of the stimulated whisker. We generated 1,800 of such structurally and functionally plausible synaptic input patterns for each of the 24 major facial whiskers, including those that correspond to the manipulated and principal whisker of our *in vivo* experiments **(Fig. 5C)**. Finally, we converted the pyramidal tract neuron into a multi-compartmental model **(Fig. S5)**, equipped with biophysical properties at the soma, dendrites, axon initial segment, and synapses that capture the characteristic intrinsic physiology of this cell type ^27^.

**Figure 5:**
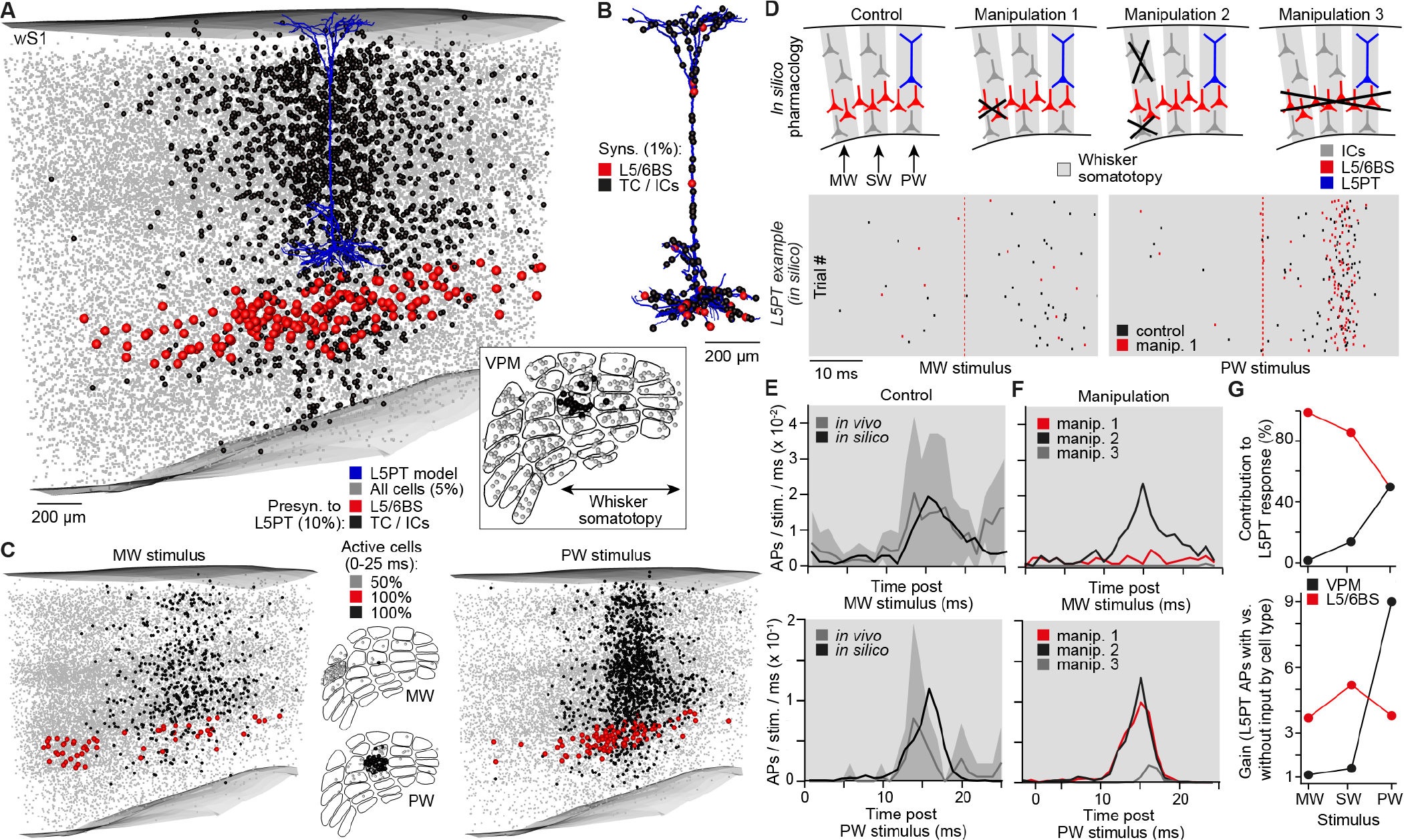
L5/6BS cells amplify and horizontally relay TC inputs to drive L5PT neurons. **A)** Example distribution of TC (i.e., inset represents model of VPM with barreloids) and IC neurons that provide synaptic input to a multi-compartmental L5PT model, embedded into an anatomically realistic model of rat wS1 ^25^. **B)** Synapse locations along the dendrites of the L5PT model corresponding to the distribution of input neurons in panel A. **C)** Example distributions of VPM and IC neurons that provide synaptic input to the L5PT model during simulations of MW or PW deflections. **D)** Dendritic integration of such generated synaptic input patterns (i.e., for deflections of the MW, SW and PW, respectively) and transformation into somatic APs were simulated for four different (pharmacology) scenarios. Raster plots represent APs of the L5PT model for 200 of these input patterns, respectively. **E)** PSTHs predicted by the L5PT model (control scenario) and as measured *in vivo* for deflections of the PW (n=9) and MW (n=8). **F)** PSTHs predicted by the L5PT model for the three different manipulation scenarios. **G)** Relative fractions of VPM and L5/6BS synapses that provided input to the L5PT model during simulations of MW, SW and PW deflections (upper panel), and the respective amplifications of L5PT AP responses (lower panel).

This multi-scale model allowed simulating how the dendrites of pyramidal tract neurons integrate and transform whisker-specific synaptic input patterns into AP output at the soma **(<u>Movie S1</u>)**. The simulations predicted responses that were indistinguishable from those recorded in pyramidal tract neurons *in vivo* **(Fig. 5D)**. In particular, AP probabilities and onsets in response to both, the principal and manipulated whisker, were in line with the respective *in vivo* data **(Fig. 5E)**. We hence performed pharmacology experiments *in silico*, at a level of spatial and cell type specificity that cannot be achieved *in vivo*. Deactivating only the border stratum cells within a volume that corresponds approximately to the spread of muscimol ^7^ abolished *in silico* responses to the manipulated whisker, but not to any other whisker **(Fig. 5F)**. Deactivating all neurons that may be affected by the muscimol, except for the border stratum cells, had in contrast no impact on the fast component of the simulated activity patterns. Deactivating all border stratum cells throughout wS1 predicted that pyramidal tract neurons lose their broadly tuned onset responses. However, the remaining direct input from the VPM is predicted to be still sufficient to evoke principal whisker responses, but with substantially reduced AP probabilities and later onsets. Our *in vivo* and *in silico* manipulations reveal that border stratum cells amplify the direct thalamocortical input of pyramidal tract neurons **(Fig. 5G)**, and relay sensory-evoked excitation from the local thalamorecipient volume to neurons across wS1, thereby driving the fast onsets of broadly tuned cortical output patterns.

We investigated which of the synaptic input parameters, or parameter combinations, of the multi-scale model could in general account for the fast onsets of sensory-evoked AP responses in pyramidal tract neurons. For each simulation trial we quantified the number of synapses that were active during the 25 ms following the stimulus onset, their respective path length distances to the soma, and times of activation at 1 ms resolution. A principal component analysis of these synaptic input statistics revealed that simulation trials with and without fast AP responses formed systematically different distributions with respect to PC_1_ **(Fig. 6A)**. 92% of the separation along the dimension of PC1 could be attributed to a single quantity **(Fig. 6B)**, in the following referred to as synchronous proximal drive (SPD). SPD represents two effective parameters: the number and synchrony of active excitatory synapses that impinge onto the proximal dendrites (i.e., path length distance to the soma < 500 μm). SPD was an almost perfect predictor **(Fig. 6C)** for AP responses (i.e., area under the receiver operating curve (AUROC) equals 1) during simulations of passive whisker deflections (AUROC = 0.83±0.03). Simulations in which we systematically varied these two parameters hence provided general relationships between the number of proximal inputs that are active within a certain time window and the resultant probability of AP responses in pyramidal tract neurons **(Fig. 6D)**. Supporting the manipulation results, these simulations predicted that– for the present experimental conditions – only combined input from the VPM and border stratum cells would be sufficiently numerous and synchronous to drive the fast APs at response probabilities which match with those observed for PW, SW and MW deflections *in vivo*.

**Figure 6:**
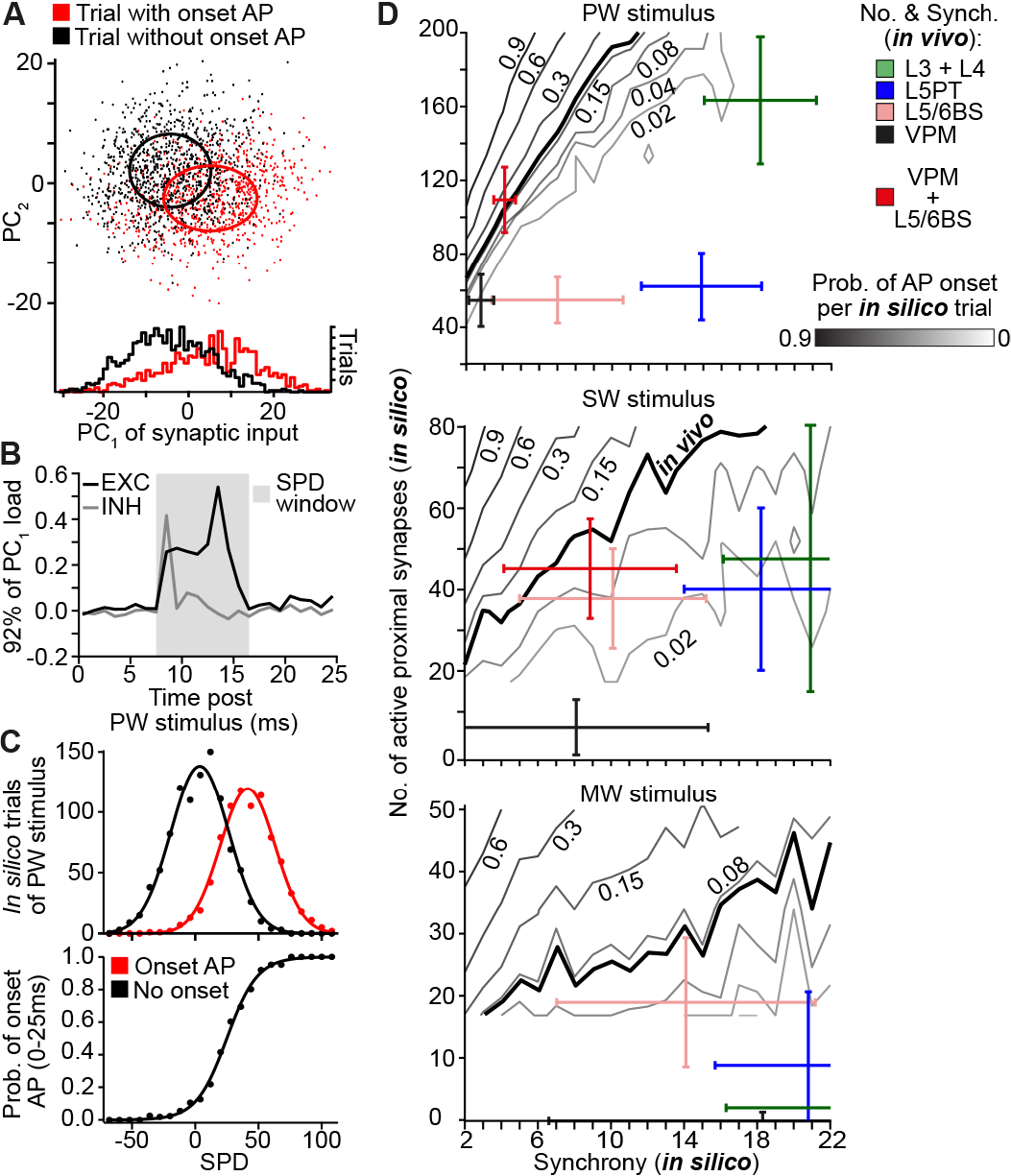
L5/6BS cells provide synchronous drive to proximal dendrites of L5PT neurons. **A.** PC analysis of spatiotemporal synaptic inputs that impinge onto the L5PT model during simulations of PW deflections. **B.** 92% of the separation between simulation trials with and without fast AP responses are reflected by PC1, which represents the net excitatory input to proximal dendrites within a time window of 8-16 ms. These two parameters are combined into a single quantity: synchronous proximal drive (SPD). **C.** SPD is an almost perfect predictor for AP responses during simulations of the L5PT model. **D.** Probabilities of fast AP responses after simulations of PW, SW and MW deflections as predicted for different combinations of the number and synchrony of proximal inputs vs. cell type-specific data derived from empirical constraints of the model. Bold lines represent our *in vivo* measurements.

## Discussion

We provide several lines of structural, functional, and computational evidence, which reveal the logic of bistratified thalamocortical input to rat wS1: preceding the vertical stream of sensory-evoked signal flow from layer 4 to layers 2/3, the strategically placed border stratum cells give rise to a second stream of excitation that spreads horizontally across layers 5/6. Parallel activation of layer 4 and a deep input stratum is likely to generalize to other sensory systems and species. For example in macaque V1, neurons – sometimes referred to as Meynert cells – have been described whose features are reminiscent of those that characterize the border stratum cells: they cluster around the layer 5/6 border and have extensive horizontal axons that span across the deep layers ^4^. The function of Meynert cells remains unknown ^28^. However, because of strong similarities in receptive field shapes between neurons in layers 4 and 6, it was suggested that these cells might be strategically placed to receive thalamocortical input from the deep layer terminal fields of lateral geniculate nucleus (LGN) axons ^4^. It was even speculated that strong thalamocortical input to horizontally projecting neurons in the deep layers represents an organizational principle that is unique to primates, and which may underlie their superior cognitive capabilities ^28^. However, bistratified LGN axons, as well as horizontally projecting thalamorecipient corticocortical neurons in the deep layers were also reported for V1 in cats ^5, 29^ and rodents ^16^.

We showed that the horizontal stream of excitation is necessary – and can even be sufficient – to drive fast sensory-evoked APs in pyramidal tract neurons. Bypassing the intracortical circuitry of the upper layers, the deep input stratum hence allows pyramidal tract neurons to integrate and transform sensory inputs from differently tuned thalamocortical populations into cortical output, which can thereby contain the entire stimulus information that was simultaneously provided by the thalamus (e.g. multi-whisker ^30^ or binocular ^31^ stimuli in wS1 or V1). In addition to providing subcortical circuits with such an integrated efference copy of the sensory input ^32^, the fast activation of pyramidal tract neurons will also be critical for intracortical computations. Somatic APs back-propagate into the apical dendrites, triggering the activation of calcium channels ^33^ that widen the pyramidal tract neurons’ time window for synaptic integration ^34^. The fast back-propagating APs that are driven by the border stratum cells will therefore switch the dendrites of pyramidal tract neurons into an active state, which occurs near simultaneous with responses in layer 4 that are driven directly by the thalamus. The two input strata could hence complement each other, ensuring that pyramidal tract neurons are able to reliably transform inputs from recurrent intracortical circuits – e.g. those from layers 2/3 that are driven by layer 4 – into cortical output ^35^. This theory is not only in line with the recent observation that sensory-evoked calcium transients in apical dendrites of pyramidal tract neurons correlate with perceptual thresholds during whisker-based behaviors ^36^. It further provides a potential explanation for the origin of sustained AP responses in pyramidal tract neurons ^14^ that persist for the duration of the stimulus **(Fig. S6)**.

The deep input stratum will be involved in other functions, beyond regulating cortical output patterns. Axons of border stratum cells innervate all layers of wS1, but in particular layer 4. The fast and reliable activation of these neurons may therefore contribute to the substantial intracortical component of sensory-evoked post-synaptic potentials in the major thalamorecipient layer ^37^. Moreover, at least a subset of the border stratum cells display long-range intrinsic axons that innervate higher-order cortices ^38^, a property that they share also with the Meynert cells ^39^. Revealing how activity patterns can be coordinated across intracortical and subcortical circuits, the parallel strata principle provides insight that will be essential for understanding how the neocortex orchestrates sensory-guided behaviors.

## Acknowledgements

We thank Bert Sakmann for discussions; Etay Hay and Idan Segev for providing biophysical models and optimization routines; Martin Schwarz for providing the AAV; Idan Segev, David Fitzpatrick and Kevan Martin for comments on the manuscript. Funding was provided by the European Research Council under the European Union’s Horizon 2020 research and innovation program (No 633428), and in part by the German Federal Ministry of Education and Research Grants BMBF/FKZ 01GQ1002 and 01IS18052, and the Deutsche Forschungsgemeinschaft (SFB 1089). **We declare that we have no conflicting interests.**

## Author contributions

M.O. conceived and designed the study. R.E. developed the model and performed simulations. R.N. performed cell-attached recordings, pharmacology experiments, and morphological reconstructions. D.U. developed analysis and data acquisition routines. A.B. performed simulations. J.G. performed virus injections and cell-attached recordings. S.D. performed extracellular recordings. C.K. performed cell-attached and extracellular recordings. All authors analyzed data. M.O. wrote the paper.

## Methods

### Animal preparation

All experiments were carried out after evaluation by the local German authorities, in accordance with the animal welfare guidelines of the Max Planck Society, or with the Dutch law after evaluation by a local ethical committee at the VU University Amsterdam, The Netherlands.

### Virus injection

Male Wistar rats aged 22-25 days (P22-25, m, Charles River) were anesthetized with isoflurane supplemented by rimadyl (Caprofen, 5mg/ kg) as analgesia, then placed into a stereotaxic frame (Kopf Instruments, model 1900), and provided with a continuous flow of isoflurane/O_2_ gas. Body temperature was maintained at 37°C by a heating pad. A small craniotomy was made above the left hemisphere 2.85 mm posterior to bregma and 3.2 mm lateral from the midline. The head of the rat was leveled with a precision of 1 μm in both the medial-lateral and anterior-posterior planes using an eLeVeLeR electronic leveling device (Sigmann Electronics, Hüffenhardt, Germany) mounted to an adapter of the stereotaxic frame. An injecting pipette containing an adeno-associated virus (AAV) was lowered into the VPM thalamus (5.05 mm from the pia). The virus – rAAV2/1-CAG-hChR2(H134R)-Syn-mCherry (titer: 1×1012 gc ml^−1^) – was provided by Martin Schwarz (University of Bonn, Germany). 50-70 nL of the virus were injected using a 30cc syringe coupled to a calibrated glass injection capillary.

### Cell-attached recording/labeling in virus injected animals

After a 16-21 day incubation period, AAV injected rats were anesthetized with urethane (1.8 g/kg body weight) by intraperitoneal injection. The depth of anesthesia was assessed by monitoring pinch withdrawal, eyelid reflexes, and vibrissae movements. Body temperature was maintained at 37.5 ± 0.5 °C by a heating pad. Cell-attached recording and labeling was performed as described in detail previously ^40^. Briefly, APs were recorded using an extracellular loose patch amplifier (ELC-01X, npi electronic GmbH), and digitized using a CED power1401 data acquisition board (CED, Cambridge Electronic Design, Cambridge, UK). APs were recorded before and during 20-30 trials of caudal multi-whisker deflections by a 700 ms airpuff (10 PSI), delivered through a 1 mm plastic tube from a distance of 8-10 cm from the whisker pad ^14^. Stimulation was repeated at constant intervals (0.3 Hz). Optical stimulation of ChR2-expressing thalamocortical terminals was provided by a 200 μm diameter optical fiber (ThorLabs #RJPSF2) coupled to a 470 nm wavelength LED (ThorLabs M470F3), resulting in an output power of 1 mW. The fiber was positioned approximately 2 mm above the cortical surface, resulting in a 1-2 mm disc of light above wS1. APs were recorded during 20-30 trials of 10 ms light pulses, at an inter-stimulus interval of 2.5 s. Following the electrophysiological measurements, neurons were filled with biocytin. Filling sessions were repeated several times. After 1-2 hours for tracer diffusion, animals were transcardially perfused with 0.9% saline followed by 4% paraformaldehyde (PFA). Brains were removed and post-fixed with 4% PFA for 24 hours, transferred to 0.05 M phosphate buffer (PB) and stored at 4°C.

### Pharmacological manipulation

Wistar rats (P28-P35, m, Charles River) were anesthetized with urethane (1.6-1.7 g/kg body weight) by intraperitoneal injection. As described above, the depth of anesthesia was monitored, and the animal’s body temperature was maintained. An ‘L’ shaped craniotomy centered on the coordinate of the barrel column representing the D2 whisker (2.5 mm posterior and 5.5 mm lateral to the bregma) was made without cutting the dura, and extended along the rostro-medial (i.e., along the E-row) and caudal axes (i.e., arc 2) for ~1-2 mm, respectively. Locations for muscimol injections and recordings were determined with long-tapered ‘search pipettes’ (tip diameter <3 μm and insertion diameter <50μm). The search pipette was inserted rostral to wS1 and lowered parallel to the midline while measuring LFPs at different cortical depths, and in response to deflections of different individual whiskers using a piezoelectric bimorph ^24^. Recordings were made using an Axoclamp 2B amplifier (Axon instruments, Union City, CA, USA), low pass filtered (300 Hz), and digitized using a CED power1401 data acquisition board (CED, Cambridge Electronic Design, Cambridge, UK). Using the LFP data, we identified the depth of the L5/6 border and the principal whisker (e.g. E2), and marked this location on the dura with a surgical pen. Repeating the LFP-guided whisker mapping with a second search pipette that was inserted approximately parallel to the vertical axis of wS1, we identified layer 5 of the hence appropriate recording site (i.e., C2 if the injection pipette was located at E2). This location was also marked on the dura. Pipettes for muscimol injections were prepared with a tip diameter of 8-12 μm. The taper diameter at the insertion point into the brain was ~125-150 μm. The tip of the pipette was filled with normal rat ringer (NRR) to avoid muscimol spill upon pipette insertion. The rest of the pipette was filled with 10 mM muscimol supplemented with 2% biocytin. The injection pipette was positioned at the previously determined location, the dura was cut open (~500 μm), and the injection pipette was inserted with positive pressure of 5-10 mbar. Allowing the tissue to adjust for 10-15 minutes, we inserted a recording pipette (i.e., 1 μm tip diameter, filled with NRR supplemented with 2% biocytin) at the second previously determined location. Both locations were confirmed by measuring whisker-evoked LFPs. Pyramidal tract neurons were identified as follows ^10, 13, 22^: (1) recording depth between 1000-1600 μm; (2) ongoing AP rates between ~1-5 Hz; (3) reliable and fast APs (i.e., between 10-20 ms) in response to principal whisker deflections; (4) reliable and fast APs after deflection of the manipulated whisker. We identified eight neurons that matched these criteria (recording / injection location: 1x B1/D1, 4x C1/E1, 3x C2/E2). Whisker deflections of the PW (e.g. C2), one SW (e.g. D2) and the MW (e.g. E2) were performed (i.e., 50 trials of 200 ms ramp-and-hold stimulus with an amplitude of ~5°, 2 s inter-stimulus interval), and APs were recorded, while simultaneously measuring the LFP via the injection pipette. Following these measurements (i.e., control data), muscimol was injected by slowly increasing the pressure onto the injection pipette (80-300 mbar), while monitoring the LFP in response to MW deflections. Once MW-evoked LFPs were abolished, and the AP activity remained unaffected, the measurements of whisker-evoked responses were repeated (i.e., at least 50 trials of PW, SW and MW deflections, respectively).

### Extracellular recordings

Wistar rats (P33-P70, m) were anesthetized using 1.6 % isoflurane in 0.4 l/h O_2_ + 0.7 l/h NO_2_, supplemented by rimadyl (Caprofen, 5mg/ kg) as analgesia. A craniotomy of 0.5 mm x 0.5 mm was made above wS1 on the left hemisphere, and a head post for fixation was implanted on the skull. After recovery from surgery, rats were head-fixed two times per day for 2-3 days. Rats quickly adjusted to the head-fixation, allowing stable recording conditions without the need of body restraint. Rats were anaesthetized with isoflurane (1.25% in 0.4 l/h O_2_ + 0.7 l/h NO_2_), and a 32-channel linear silicon probe (E32+R-50-S1-L10(NT), Atlas Neuroengineering, Belgium) was inserted into wS1 for extracellular multi-unit recordings. Prior to recordings, silicon probes were labeled with DiI (Thermo Fisher Scientific, Waltham, MA, USA). The probe was connected to a unity-gain headstage (Neuralynx, USA), in series with the Open Ephys data acquisition board equipped with a RHD2132 digital interface chip (Intan Technologies, Los Angeles, CA, USA). Using the LFP strategy described above, the PW at the recording site was identified, all other whiskers were trimmed to 5 mm, and the anesthesia was terminated. Recordings were performed once the animals were fully awake (~25 minutes after the anesthesia was terminated ^41^). Rats were not trained to perform tactile behavior, and behavior was not rewarded. Sensory input resulted from whisker touch with a pole that was placed within range during periods of exploratory whisker self-motion. The touch onset was determined by high-speed videography at 200 frames/s (MotionScope M3 camera, IDT Europe, Belgium). Whisker angle was tracked offline ^42^, and episodes of whisker movements were classified by thresholding average power in whisker angle versus time (1-20 Hz bandpass) using the Matlab spectrogram function. Touch events were detected manually in each frame. Signals were acquired at a sampling rate of 30 kHz/channel using Open Ephys GUI ^43^. To identify single units, the data were high-pass filtered, and automatically sorted into clusters using Klustakwik ^44^. The clusters were manually post-processed, and only stable and well-isolated single units were considered for analysis. The average waveforms of all well-isolated single units were used to sub-classify units ^45^ as regular spiking vs. fast spiking units (FSUs). FSUs (AP peak-to-trough time <0.5 ms and AP half-peak time <0.25 ms) were excluded from the analyses. After recordings, rats were anaesthetized with urethane (>2.0 g/kg) and perfused with 0.9% NaCl followed by 4% paraformaldehyde (PFA).

### Histology

For morphological reconstructions, 100 μm thick vibratome sections were cut tangentially to wS1 (45° angle) ranging from the pial surface to the white matter (WM). Sections were processed for cytochrome-C oxidase staining to visualize barrel contours in layer 4 ^46^. All sections were treated with avidin-biotin (ABC) solution, and subsequently neurons were identified using the chromogen 3,3′-diaminobenzidine tetrahydrochloride (DAB) ^47^. All sections were mounted on glass slides, embedded with Mowiol, and enclosed with a cover slip. In experiments where AAV injections were combined with biocytin filling, cortex was cut into 45-48 consecutive 50 μm thick tangential sections. Sections were treated with Streptavidin Alexa-488 conjugate (5mg/ml Molecular Probes #S11223) to stain biocytin labeled morphologies ^14^. To enhance the virus labeling, sections were immunolabeled with anti-mCherry antibody. Sections were permeabilized and blocked in 0.5% Triton x-100 (TX) (Sigma Aldrich #9002-93-1) in 100 mM PB containing 4% normal goat serum (NGS) (Jackson ImmunoResearch Laboratories #005-000-121) for 2 hours at room temperature. The primary antibody was diluted 1:500 (Rabbit anti-mCherry, Invitrogen #PA5-34974) in PB containing 1% NGS for 24 hours at 4°C. The secondary antibody was diluted (1:500 goat anti-Rabbit IgG Alexa-647 H+L Invitrogen #A21245) and was incubated for 2-3 hours at room temperature in PB containing 3% NGS and 0.3% TX. All sections were mounted on glass slides, embedded with SlowFade Gold (Invitrogen #S36936) and enclosed with a cover slip. For extracellular recording experiments, brains were post-fixed in 4% PFA, and tangential vibratome sections (100 μm) were cut and stained for cytochrome-C. An X-Cite 120 Q light-source (Excelitas Technologies Corp., Waltham, MA, USA) was used to visualize the DiI electrode tract, and only electrode tracks within the barrel column that represents the PW were selected for analyses. The histology allowed assigning the recording depth to each electrode (i.e., and hence to each unit) with approximately 100 μm precision.

### Morphological reconstruction

Neuronal structures were extracted from image stacks using a previously reported automated tracing software ^48^. 3D image stacks of up to 5 mm × 5 mm × 0.1 mm were acquired using an automated brightfield microscope system (BX-51, Olympus, Japan) at a resolution of 0.092 × 0.092 × 0.5 μm per voxel (100× magnification, NA 1.4). For reconstruction of fluorescently labeled neurons, images were acquired using a confocal laser scanning system (Leica Application Suite Advanced Fluorescence SP5; Leica Microsystems). 3D image stacks of up to 2.5 mm × 2.5 mm × 0.05 mm were acquired at a resolution of 0.092 × 0.092 × 0.5 μm per voxel (63× magnification, NA 1.3). Manual proof-editing of individual sections, and automated alignment across sections were performed using custom-designed software ^49^. Pia, barrel and WM outlines were manually drawn on low resolution images (4×). Using these anatomical reference structures, all reconstructed dendrite and axon morphologies were registered to the D2 barrel column of a standardized 3D reference frame of rat wS1^9^. The shortest distance from the pial surface to the soma, and 20 morphological features that have previously been shown to separate between excitatory cell types in rat wS1 ^9^ were calculated for each reconstructed and registered dendrite morphology. For identification of putative thalamocortical synapses, biocytin labeled morphologies and AAV labeled VPM terminals were imaged simultaneously using the confocal laser scanning system as described above: biocytin Alexa-488 (excited at 488 nm, emission detection range 495-550 nm), AAV Alexa-647 (excited at 633 nm, emission detection range 650-785 nm). These dual-channel 3D image stacks were loaded into Amira visualization software (FEI). All reconstructed dendrites were manually inspected, and landmarks were placed onto each spine head. If a spine head was overlapping with a VPM bouton, an additional landmark was placed to mark a putative synapse. The shortest distance of each landmark to the dendrite reconstruction was determined, and the path length distance was calculated from that location along the reconstructed L5/6BS cell to the soma.

### Cell type-specific analysis

In total, n=177 *in vivo* labeled morphologies of excitatory neurons in wS1 (i.e., from urethane anesthetized Wistar rats; P25-P45, m/f, Charles River) were used in this study to determine cell type-specific whisker receptive fields (wRFs), and to provide structural/functional constrains for simulation experiments. All morphologies ^12, 14^ – except for five L5/6BS and one L5PT neurons – and classification approaches ^12, 13^, as well as the corresponding whisker-evoked physiology data ^10, 13^ have been reported previously, but in different context. Analysis of wRFs for objectively classified morphological cell types were not performed for any of the previously reported neurons. Here, each neuron was objectively assigned to one of the 10 major excitatory cell types of the neocortex ^2, 12^ based on the 21 soma-dendritic features described above: three types of pyramids in layers 2-4 (L2PY, L3PY, L4PY), spiny-stellates (L4ss) and star-pyramids in layer 4 (L4sp), slender-tufted intratelencephalic (L5IT) and thick-tufted pyramidal tract neurons in layer 5 (L5PTs), polymorphic corticocortical (L6CC) and corticothalamic neurons in layer 6 (L6CT), and the L5/6 border stratum cells (L5/6BS). In the present study, we grouped L4ss and L4sp as layer 4 spiny neurons (L4SP). The physiology data (i.e., AP responses to passive deflections of the principal and its eight adjacent whiskers ^10^) were grouped by the hence determined morphological cell types, resulting cell type-specific wRFs.

### Multi-compartmental model

We generated a biophysically-detailed multi-compartmental neuron model, which captures the stereotypic morphological and intrinsic physiological properties of L5PTs. The L5PT model is based on the 3D soma-dendrite reconstruction of a L5PT neuron, whose morphological and topological features – which allow discriminating L5PTs from other excitatory cell types in the deep layers (see above) – represent approximately the respective averages across a population of 37 L5PTs ^12, 14^ that were labeled *in vivo* via cell-attached recordings in layer 5 of rat wS1. A simplified axon morphology was attached to the reconstructed soma based on ^50^. The axon consisted of an axon hillock with a diameter tapering from 3.5 μm to 1 μm over a length of 10 μm, an axon initial segment (AIS) of 10 μm length and 1 μm diameter, and 1 mm of myelinated axon (diameter of 1 μm). The diameter of the reconstruction of the apical trunk and oblique dendrites was scaled by a factor of 2.5 to allow for backpropagation of action potentials (bAP), and bAP-triggered calcium spike (BAC) firing to occur (i.e., after scaling the diameter of the apical trunk was 4.5 μm at the soma, and 1.5 μm at the main bifurcation located at a distance of ~900 μm from the soma). Spatial discretization of the dendrite morphology (i.e., compartmentalization) was performed by computing the electrotonic length constant of each dendrite branch at a frequency of 100 Hz and setting the length of individual compartments in this branch to 10% of this length constant. The length of axonal compartments was set to 10 μm. After spatial discretization, the L5PT morphology consisted of 1033 compartments with an average length of ~15 μm, but no longer than 42 μm. The resultant L5PT model was then combined with previously reported biophysical models of a variety of Hodgkin-Huxley (HH)-type ion channels **(Table S1)** that are expressed at different densities within the soma, basal and/or apical dendrites, and axon initial segment ^27^. Using an evolutionary multi-objective optimization algorithm ^51^, we tuned the parameters of the biophysical models until numerical simulations of the L5PT model (using NEURON 7.2 ^52^) reproduced current injection-evoked somatic and/or dendritic sub- and suprathreshold responses that are characteristic for L5PTs, as measured previously via whole-cell recordings in acute brain slices of rat wS1 *in vitro* ^27^. Fixed membrane parameters were the axial resistance (100 Ωcm in all compartments), the membrane capacitance (1 μF/cm^2^ at the soma and axon, 2 μF/cm^2^ in the apical and basal dendrites to account for increased surface area due to spines, and 0.04 μF/cm^2^ along the myelinated part of the model axon), and the passive membrane conductance along the myelinated part of the axon (g_pas_ = 0.4 pS/μm^2^, i.e., equivalent to a specific membrane resistance of 25,000 Ωcm^2^). The reversal potential of the passive membrane conductance was set to −90 mV. Conductance densities of the non-specific cation current I_h_ were fixed at 0.8 pS/μm^2^ in the soma and axon, and 2 pS/μm^2^ in the basal dendrites. In the apical dendrite, the conductance density of I_h_ increased exponentially with the distance to the soma. The biophysical model parameters to be optimized were the peak conductance per unit membrane area for various voltage-dependent ion channels, and the parameters of a phenomenological model of the calcium dynamics in different parts of the morphology (i.e., axon, soma, basal and apical dendrites; **Table S1**). The targets of the optimization were different features of the membrane potential in response to two stimuli, as measured previously ^27^: (1) a brief current injection into the soma should trigger an AP at the soma and a bAP, and (2) a brief current injection into the soma, followed by current injection into a Ca^2+^ channel dense region around the first bifurcation point of the apical tuft, should trigger somatic bursts (i.e., BAC firing). The specific features, as listed in **Table S2**, were combined into five objectives, which were then optimized simultaneously by using the evolutionary algorithm ^51^. A set of 1,000 models was generated with parameters drawn randomly from a physiologically plausible range. In every iteration, each model was then evaluated by simulating the response to the two stimuli, calculating the features and determining the error by calculating the difference between each simulated and measured feature in units of standard deviations of the experimental feature ^27^. After each model had been evaluated, a new set of 1,000 models was generated from the previous set by stochastically transferring parameter values from “good” models (i.e., lower errors) to “worse” models (i.e., higher errors). Additionally, parameter values of all models were updated stochastically to avoid converging to local minima. This procedure was repeated 500 times. From the final iteration, the set of biophysical models used here was selected based on three criteria: (1) it had the lowest sum across all objective errors, (2) similar deviations in all objective errors (i.e., models where only a subset of objectives matched the experimental data were not considered), and (3) it supported regular spiking of increasing frequencies in response to sustained current injections of increasing amplitude.

### Connectivity model

The structurally plausible constraints for the numbers and dendritic distributions of cell type-specific synaptic input patterns that impinge onto the L5PT model are based on an anatomically realistic network model of rat wS1, as described in detail previously ^25^. Briefly, we generated a 3D model of the average geometry of rat wS1 (i.e., 3D location, orientation and diameter of all barrel columns; 3D pial and white matter (WM) surfaces), and determined the variability (~50 μm) of these anatomical landmarks across twelve animals ^9^. Next, we measured the number and 3D distribution of all excitatory and inhibitory neuron somata in rat wS1 (~530,000 neurons) and the ventral posterior medial nucleus (VPM) of the thalamus (~6,000 neurons) in four different animals ^11^, and generated an average excitatory and inhibitory 3D neuron somata distribution at a resolution of 50×50×50 μm^3^, reflecting the variability of the cortex geometry across animals. We then registered a sample of 177 excitatory intracortical (IC) neuron morphologies (i.e., grouped into ten cell types (see above) ^10, 12^, 14 excitatory thalamocortical (TC) axon morphologies labeled in VPM *in vivo* ^6^, and the soma-dendrites of 213 inhibitory neuron (IN) morphologies (203 labeled in L2-6 *in vitro* ^53, 54, 55^, 10 labeled in L1 *in vivo* ^56^) to the geometric model of wS1. Combining these data by using a previously reported network building approach ^25^, we generated a structurally dense model of wS1, which comprised soma, dendrite and axon morphologies that represent all of the excitatory (here: 462,402) and inhibitory neurons (here: 67,535) that are located in rat wS1, as well as axon morphologies that represent the IC part of all VPM neurons (here: 6,225). To estimate synaptic connectivity within this structurally dense wS1 model, we calculated the overlap at 50 μm^3^ resolution between the putative postsynaptic target structures (PSTs; i.e., soma/dendrite surface for inhibitory connections; dendritic spines for excitatory connections) and putative presynaptic sites (i.e., axonal boutons) for all pairs of neurons, and normalized this quantity by the respective total amount of locally available PSTs (i.e., total somatic/dendritic surface and number of spines within each 50 μm voxel). Neglecting wiring specificity at subcellular scales ^25^, we converted these overlap measurements into connection probabilities, which predict the respective distributions of the numbers and most likely dendritic locations of synaptic contacts. To compare the predicted connection probabilities between excitatory IC cell types and L5PTs with previously reported paired-recording results that were obtained from acute brain slices *in vitro*, we cropped out ten 300 μm wide thalamocortical/semi-coronal slices from the network model, which comprised at least half of the C2 barrel column volume. Connection probabilities that were predicted for truncated morphologies in slices are denoted by asterisks in **Table S3**. To compare the predicted connection probabilities between TC neurons and L5PTs with previously reported paired-recording results that were obtained *in vivo*, we used L5PTs whose somata were closest to the C2 barrel column (i.e., including septal neurons) and TC neurons located in the C2 VPM-barreloid of the uncropped network model. To compare the predicted connection probabilities between INs and L5PTs with previously reported paired-recording results we grouped all excitatory neurons in layer 5 (i.e., L5PTs and L5ITs) whose somata were closest to the C2 barrel column (i.e., including septal neurons) and INs whose axons remained largely confined to layer 5. Finally, we embedded the L5PT model into the network model of wS1 by using a previously reported registration approach ^9^. Here, we placed the L5PT model at nine different locations within the barrel column representing the C2 whisker (i.e., approximately in the center of wS1), while preserving its (*in vivo*) soma depth location. For each of the nine locations (i.e., one in the column center, and eight at equally spaced angular intervals with a distance of ~100 μm to the column center) we used the connectivity mapping procedures as described above to estimate the numbers and dendritic locations of cell type-specific synaptic inputs that impinge onto the dendrites of the L5PT model. Specifically, by sampling from the overlap distributions 50 times, calculating the mean of the number of synaptic inputs from each cell type, and choosing the sample that was closest to this mean, we estimated that the L5PT model receives a total of 24,161 ± 785 synaptic inputs. Of those, ~90% are predicted to originate from excitatory IC and TC neurons, which corresponds to an average density of 1.4 glutamatergic and 0.14 GABAergic synapses per μm dendrite, respectively (i.e., 148 ± 18 GABAergic synapses are located on the soma).

### Synapse models

Conductance-based synapses were modeled with a double-exponential time course. Excitatory synapses contained both AMPA receptors (AMPARs) and NMDARs. Inhibitory synapses contained GABAARs. The reversal potential of AMPARs and NMDARs was set to 0 mV, that of GABAARs to −75 mV. Rise and decay time constants of AMPARs were set to 0.1 ms and 2 ms, respectively ^57^; those of NMDARs to 2 ms and 26 ms, respectively 57; and those of GABAARs to 1 ms and 20 ms, respectively ^58^. The Mg-block of NMDARs was modeled by multiplying the conductance value with an additional voltage-dependent factor 1/(1 + η · exp(−γ · V)) ^59^, where η = 0.25, γ = 0.08/mV, and V is the membrane potential in mV ^60^. The peak conductance at excitatory synapses from different presynaptic cell types was determined by assigning the same peak conductance to all synapses of the same cell type, activating all connections of the same cell type (i.e., all synapses originating from the same presynaptic neurons) one at a time, and comparing parameters of the resulting unitary postsynaptic potential (uPSP) amplitude distribution (mean, median and maximum) for a fixed peak conductance with experimental measurements *in vitro* (input from L2-6 ^61^) or *in vivo* (TC input ^7^). The peak conductance for synaptic inputs from each cell type was systematically varied until the squared differences between the parameters of the *in silico* and *in vitro*/*in vivo* uPSP amplitude distributions were minimized **(Table S4)**. The peak conductance at inhibitory synapses was fixed at 1 nS ^35^. Release probability at excitatory and inhibitory synapses was fixed at 0.6 and 0.25, respectively ^35, 62^.

### Synaptic input patterns

Synaptic input patterns to the L5PT model were estimated as follows: All presynaptic neurons determined during the network-embedding procedure were converted into point neurons that could emit APs. During periods of ongoing activity, APs in presynaptic neurons were modeled as Poisson trains with cell type-specific mean firing rates as measured *in vivo* ^13^. The mean firing rate of INs was set to 7 Hz ^35^ (except for L1 INs ^56^). Each AP in a presynaptic neuron is registered at all synapses between the presynaptic neuron and the L5PT model without delay and may cause a conductance change, depending on the release probability of the synapse. After a stimulus (i.e., deflection of the PW, SW or MW), each presynaptic neuron can emit additional spikes. The location of the deflected whisker in the wRF of the presynaptic neuron is determined based on the barrel column where the soma of the presynaptic neuron is located in (i.e., a convolution operation), and the corresponding whisker-specific post-stimulus time histogram (PSTH) is used to stochastically generate additional sensory-evoked APs. Whisker-specific PSTHs of excitatory cell types were generated based on *in vivo* wRF measurements (**Fig. S3)**. The amplitude of the PSTH of excitatory IC cell types is further scaled by a factor of 0.4571 to reflect lower response probabilities of cortical neurons in the up-state ^63^. The whisker-specific PSTHs of TC neurons in VPM were constructed based on previously published *in vivo* measurements, where single- and multi-whisker responsive neurons were described for the same experimental conditions used in this study ^23^. Single- and multi-whisker responsive VPM neurons were grouped into a single TC PSTH. The whisker-specific PSTHs of INs in wS1 were constructed based on previously published *in vivo* measurements, which were acquired under the same experimental conditions that were used here ^26, 64^, and which can be summarized as follows: (1) the onset times of whisker-evoked APs in INs across all layers should be similar to those of the excitatory cell types; (2) in case of PW touch, AP onset times in INs should precede those of the excitatory IC, but not TC cell types; (3) INs have broad wRFs. To capture these empirical constraints, the PW/SW-evoked PSTHs of INs were set to the respective maximum values across all excitatory cell types in each 1 ms time bin; the resultant PW-evoked PSTH was shifted by −1 ms (but no spiking before TC neurons; i.e., > 8 ms); and the ratio between the integrals of the PW- and SW-evoked PSTHs during 0-50 ms was set to a fixed ratio of 2:1. These constraints leave one free parameter for constructing the PSTHs of INs: the total number of PW-evoked APs during 0-50 ms post stimulus. We simulated the response of the L5PT model after PW deflections while systematically varying this parameter, and computed the resulting number of APs during 0-25 ms, until the L5PT model exhibited simulation trials with and without AP responses as measured *in vivo*. This yielded a value for INs of 1.0 APs per PW deflection per 50 ms. *In silico* pharmacology experiments were performed by removing all synaptic inputs from specific presynaptic populations from the model as shown in **Fig. 5D**.

### Simulations

We generated 200 samples of structurally- and functionally-plausible cellular stimulus representations for each of the nine L5PT model locations (i.e., 1,800 samples per whisker), and for each simulated whisker deflection in the control condition (i.e., the complete network model), and for three different *in silico* pharmacology experiments. Since the L5PT model was located in the C2 column, simulated C2 deflections were assigned as PW deflections, those of the eight adjacent whiskers as SW deflections, and simulated E2 deflections as MW deflections. The three different *in silico* pharmacology conditions were as follows: (1) synapses from border stratum (L5/6BS) cells whose somata were located within the E2 column or the surrounding septum were removed from the L5PT model; (2) synapses from neurons of all excitatory cell types whose somata where located within the E2 column, except for L5/6BS cells, were removed; (3) synapses from all L5/6BS cells throughout wS1 were removed. All combinations of L5PT model location, identity of the deflected whisker, and pharmacology condition resulted in 72,000 spatiotemporal synaptic input patterns, which we associate with different trials. For each trial, we numerically simulated the integration of the respective conductance changes within all dendritic compartments (and the soma and axon) of the HH-type L5PT model. Each simulation trial consisted of 245 ms ongoing activity, followed by 50 ms of sensory-evoked activity. The first 100 ms and the last 25 ms of simulated activity were discarded. AP times were determined from zero-crossings of the simulated membrane potential at the soma. For each of the simulation trials (control condition) we created a 100-dimensional vector, which quantified the spatiotemporal features of the respective synaptic input patterns that impinge onto the L5PT model. Entries of the vector represented all active synapses during the period of 0-25 ms post stimulus, their respective path length distances to the soma, times of activation with 1 ms resolution, and whether the synapses originated from excitatory or inhibitory neurons. The input vectors were sorted into two groups, representing simulations in which onset APs (i.e., during the period of 8-25 ms post stimulus) did or did not occur. A principal component analysis (PCA) of these spatiotemporal input vectors revealed that trials with vs. without onset APs formed overlapping, but systematically different distributions. PC_1_ discriminated between these distributions. 92% of PC_1_ could be accounted for by the difference between excitatory and inhibitory inputs that are active during a period of 8-16 ms post stimulus, and that are located within less than 500 μm path length distance to the soma (here referred to as proximal inputs). We defined a single quantity that represents PC_1_ – synchronous proximal drive (SPD) – i.e., it reflects the net input (i.e., number of active excitatory minus active inhibitory synapses) along the proximal dendritic compartments of the L5PT model (i.e., path length distance <500 μm) within 8-16 ms. We then calculated the probability of observing a whisker-evoked AP response in the L5PT model as a function of SPD, and fitted a sigmoidal curve to this distribution. The inverse width (or slope) of the fitted sigmoidal curve can be interpreted as a measure for the predictive power of SPD for AP responses. We systematically varied the end time point of the integration time window to determine the SPD window with highest predictive power, which matched closely with the SPD window determined for PW deflections by the PCA. These SPD windows were then used to compute the AUROC values for PW and SW deflections reported in the main text. Breaking down SPD into its two parameter, (1) the number of active excitatory synapses along the proximal dendrites, and (2) their respective synchrony (i.e., time window in which they are active), we performed additional simulations of the L5PT model. To do so, the structurally and functionally constrained PW/SW-evoked spatiotemporal synaptic input patterns were replaced as follows. All other model parameters (i.e., biophysical and synapse models), as well as synaptic input patterns preceding the stimulus remained unchanged. First, the distribution of stimulus evoked synaptic inputs along the dendrites of L5PT model was determined by calculating the average distribution of active synapses during 50 ms following PW and SW simulation trials (i.e., from the structurally and functional constrained trails). Second, the resultant 3D distributions of active excitatory and inhibitory synapses were converted into distance-dependent probability distributions (i.e., 1D) with 50 μm (i.e., path length) resolution. Third, the subcellular distributions, temporal activation patterns and numbers of active synapses (i.e., excitatory/inhibitory during periods of ongoing activity; inhibitory during periods of whisker-evoked activity) were then determined by calculating the respective averages across PW and SW simulation trials (i.e., from the structurally and functional constrained trails), respectively. Fourth, the temporal distribution of active excitatory synapses was modeled as a log-normal distribution ^10^ with a fixed offset of 8 ms post-stimulus (i.e., corresponding to the onset latency of VPM input) and a fixed peak time of 9 ms post-stimulus. Fifth, the only remaining parameter was the median timing of the log-normal distribution. Varying this parameter in 1 ms steps resulted in excitatory synaptic input distributions that ranged from highly synchronous (2 ms; i.e., median timing at 10 ms post stimulus) toward asynchronous (i.e., median timing much later than 10 ms post stimulus). Sixth, at the same time, the total number of active excitatory synaptic inputs was systematically varied. Seventh, for each combination of the number and synchrony of stimulus evoked excitatory inputs, 200 samples of spatiotemporal synaptic input patterns were generated and simulated as described above. Then, the probability of an onset AP (i.e., between 8-16 ms) was calculated for each combination of the number and synchrony of stimulus evoked excitatory inputs. Iso-AP probability contour plots were calculated by arranging all synaptic input number and synchrony combinations in a 2D grid, and linear interpolation between the grid points. The corresponding *in vivo* data of cell type-specific numbers and synchronies of active proximal inputs were derived from the structural and functional simulation constraints of PW, SW and MW deflections (i.e., representing our *in vivo* measurements **(Fig. S3)** and reconstructions of the network model **(Fig. S5)**).

### Statistical analysis

All data are reported as mean ± standard deviation unless mentioned otherwise. Normality was not assumed when performing statistical testing. All tests were performed using the R software package (version 3.4.3) and the scipy python package (version 1.0.1).

### Data availability

All relevant data are available from the authors. The model and simulation routines, including a detailed documentation of all parameters and the analysis routines can be obtained from ModelDB (http://senselab.med.yale.edu/ModelDB/; accession number: 239145; password: Horizontal).

## Supplementary Materials

**Figure S1:**
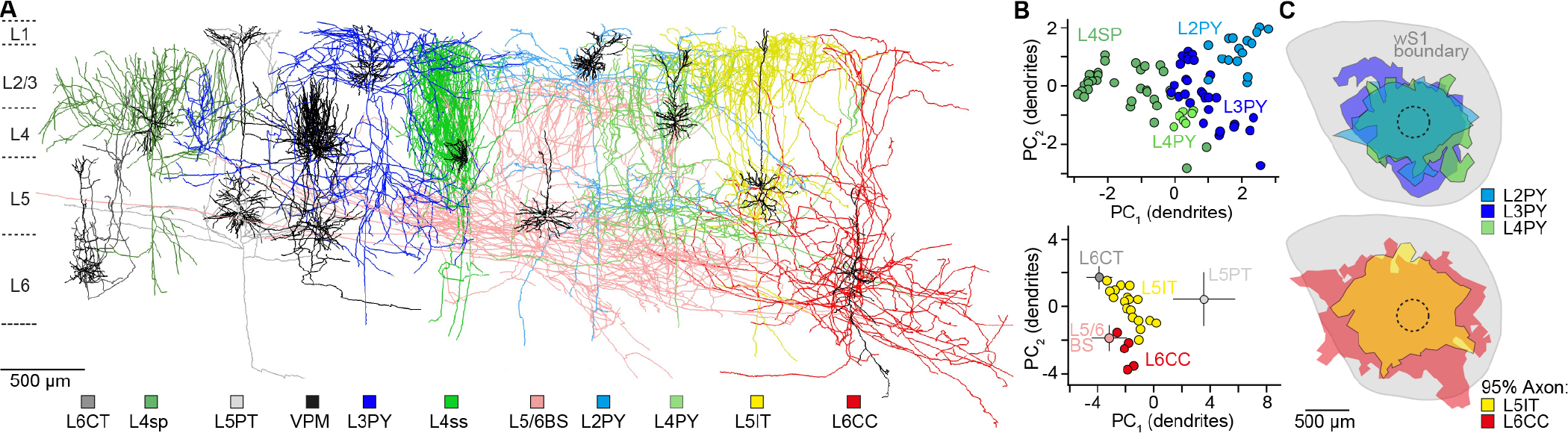
Cell type-specific structural constrains for *in silico* experiments. **A)** Gallery of exemplary *in vivo* labeled neuron morphologies for each of the 10 major excitatory cell types of the neocortex, whose soma, dendrite and axon distributions (ICs, n=177) were compared with the laminar distribution of thalamocortical axons from the ventral posterior medial nucleus (VPM, n=14). Neurons were classified as reported previously ^12^ into pyramidal neurons in layer 2 (L2PY, n=16), layer 3 (L3PY, n=30) and layer 4 (L4PY, n=7), spiny stellates (L4ss, n=22) and star-pyramids in layer 4 (L4sp, n=15), slender-tufted intratelencephalic (L5IT, n=18) and thick-tufted pyramidal tract neurons in layer 5 (L5PT, n=37), corticothalamic (L6CT, n=13) and polymorphic corticocortical neurons in layer 6 (L6CC, n=5), and corticocortical neurons at the layer 5/6 border (L5/6BS, n= 14). L4ss and L4sp neurons were grouped as layer 4 spiny neurons (L4SP). **B)** Principal components (PC_1/2_) of dendritic features that discriminate between excitatory cell types in the upper and deep layers, respectively. The means and STDs of L5/6BS, L5PT and L6CT neurons from **Fig. 1C** are shown for comparison. **D)** Horizontal axon extent of L2PY (n=9), L3PY (n=15), L4PY (n=4), L5IT (n=5) and L6CC (n=4) neurons.

**Figure S2:**
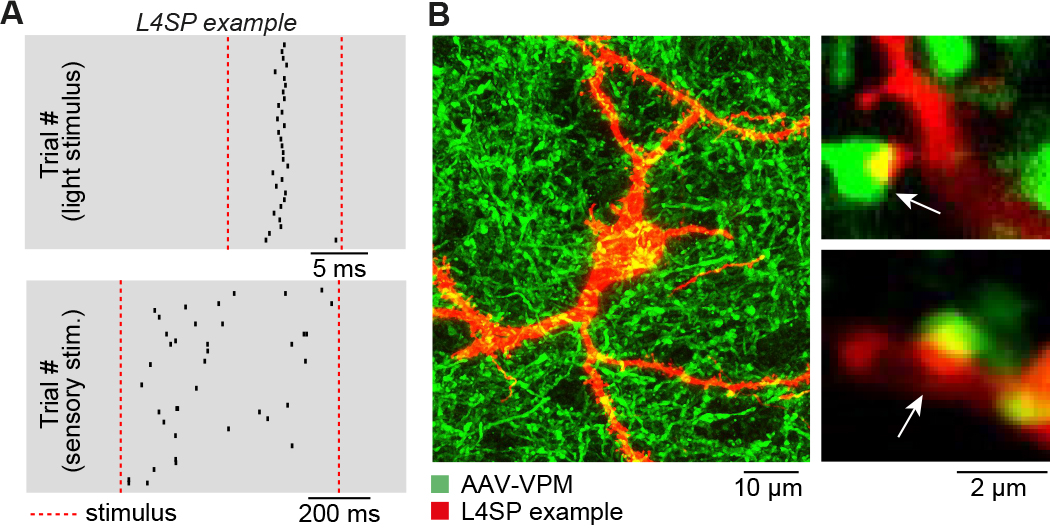
Thalamocortical input to L4SP cells. **A)** Example of cell-attached *in vivo* recording in layer 4 of wS1 of AAV-injected brain. Ticks represent APs in response to a 10 ms flash of green light onto the cortical surface (top), and a 700 ms airpuff onto the whiskers (bottom). **B)** Confocal images of the neuron shown in panel A. The neuron was morphologically identified as L4SP. Putative thalamocortical synapses were identified as contacts between VPM boutons and dendritic spines.

**Figure S3:**
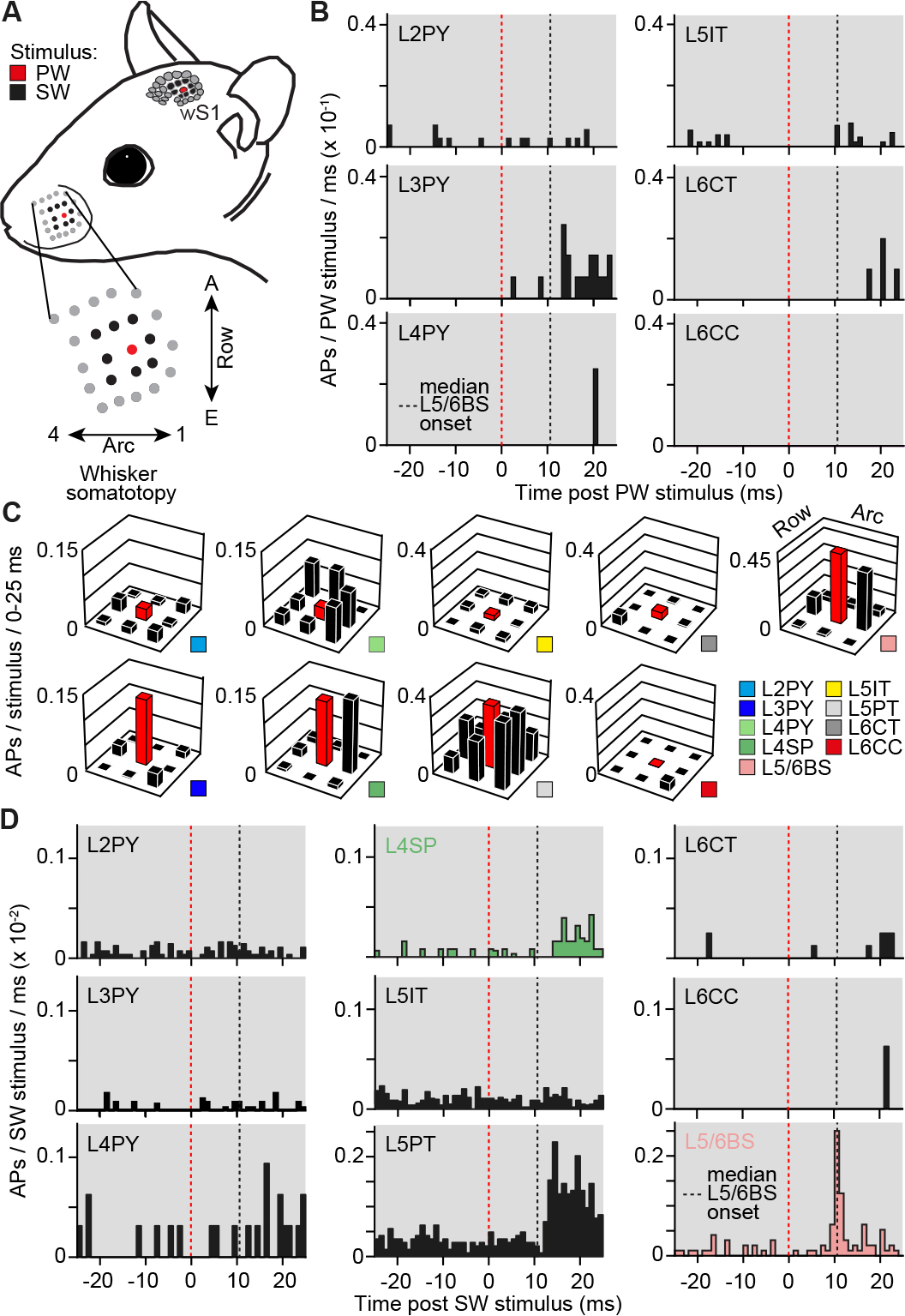
Cell type-specific functional constrains for *in silico* experiments. **A)** Illustration of cell type-specific mapping of whisker receptive fields (wRFs) as reported previously ^10^ (i.e., under conditions that were consistent with those of the pharmacological manipulations). Action potentials (APs) were recorded in responses to deflections of the principal whisker (PW), and of the eight whiskers that are adjacent to the PW (here referred to as SWs). **B)** PSTHs of PW-evoked APs for morphologically classified L2PY (n=7), L3PY (n=7), L4PY (n=2), L5IT (n=13), L6CT (n=5) and L6CC (n=1) neurons, analogous to those shown in **Fig. 3C** for L4SP, L5PT and L5/6BS neurons. **C)** Cell type-specific wRFs representing the cells in panels B and D. **D)** PSTHs of SW-evoked APs for all cell types (i.e., averaged across the adjacent whiskers), representing the cells shown in panels B and C, and **Fig. 3C**.

**Figure S4:**
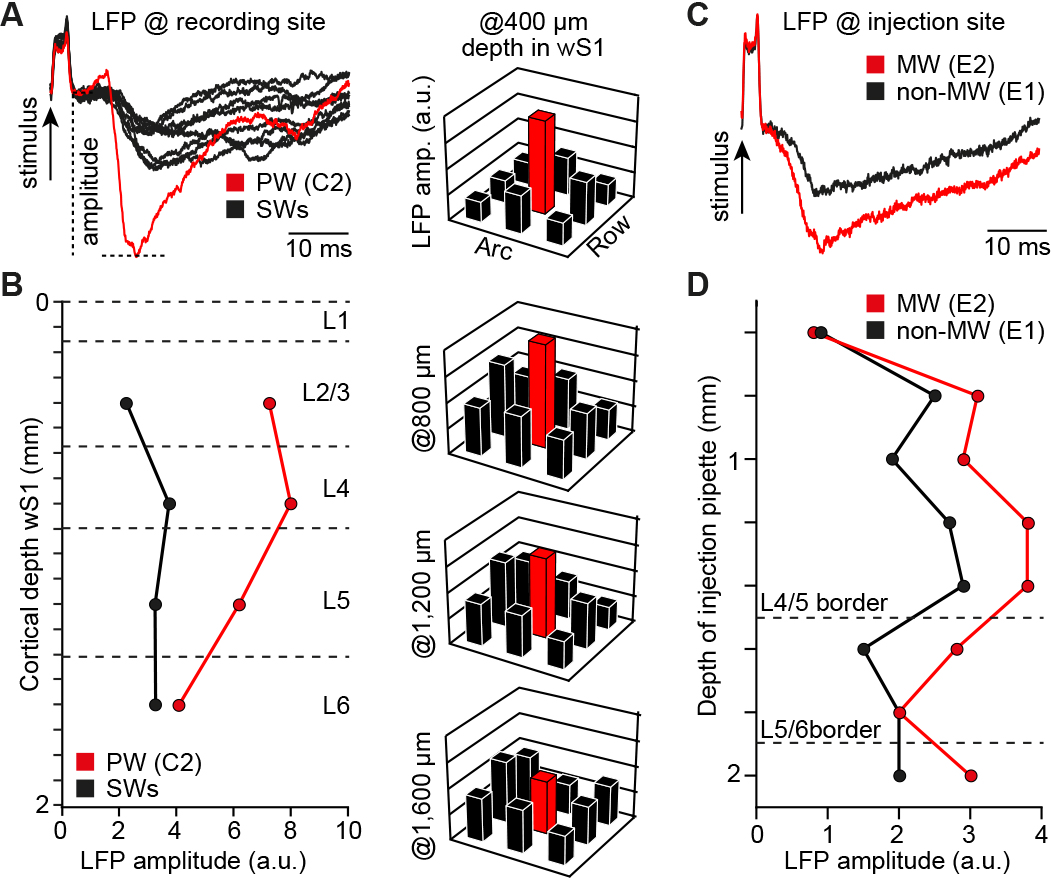
LFP guided *in vivo* pharmacology. **A)** Left panel: LFP recordings via search pipette at 400 μm depth in wS1. LFP amplitudes in response to deflections of the PW and its eight SWs were quantified. Right panel: LFP wRF reveals the PW at the recording site ^24^ (here: C2). **B.** LFP wRF measurements were repeated at different cortical depths of wS1. Using the depth of layer borders ^11^, the characteristic laminar profiles of LFP responses to PW (and SW) stimuli were used to identify the border between layer 4 and 5 (i.e., ~100 μm below the LFP maximum). The target location at the L5/6 border was hence approximately 400-500 μm below the LFP maximum. **C-D)**. Example experiment that illustrates how the LFP depth profile was used to locate the L5/6 border of the barrel column representing the MW whisker (i.e., E2). The muscimol injection pipette was inserted rostral to wS1 at an angle that was approximately parallel to the midline (i.e., oblique to the vertical axis of wS1). E2 was identified as the MW based on the larger LFP amplitudes across the cortical depth when compared to those evoked by SW stimuli (shown here: E1). The target location (i.e., L5/6 border) was then determined by identifying the depth of maximal LFP amplitude and adding 500 μm (i.e., here injection at ~1850 μm depth).

**Figure S5:**
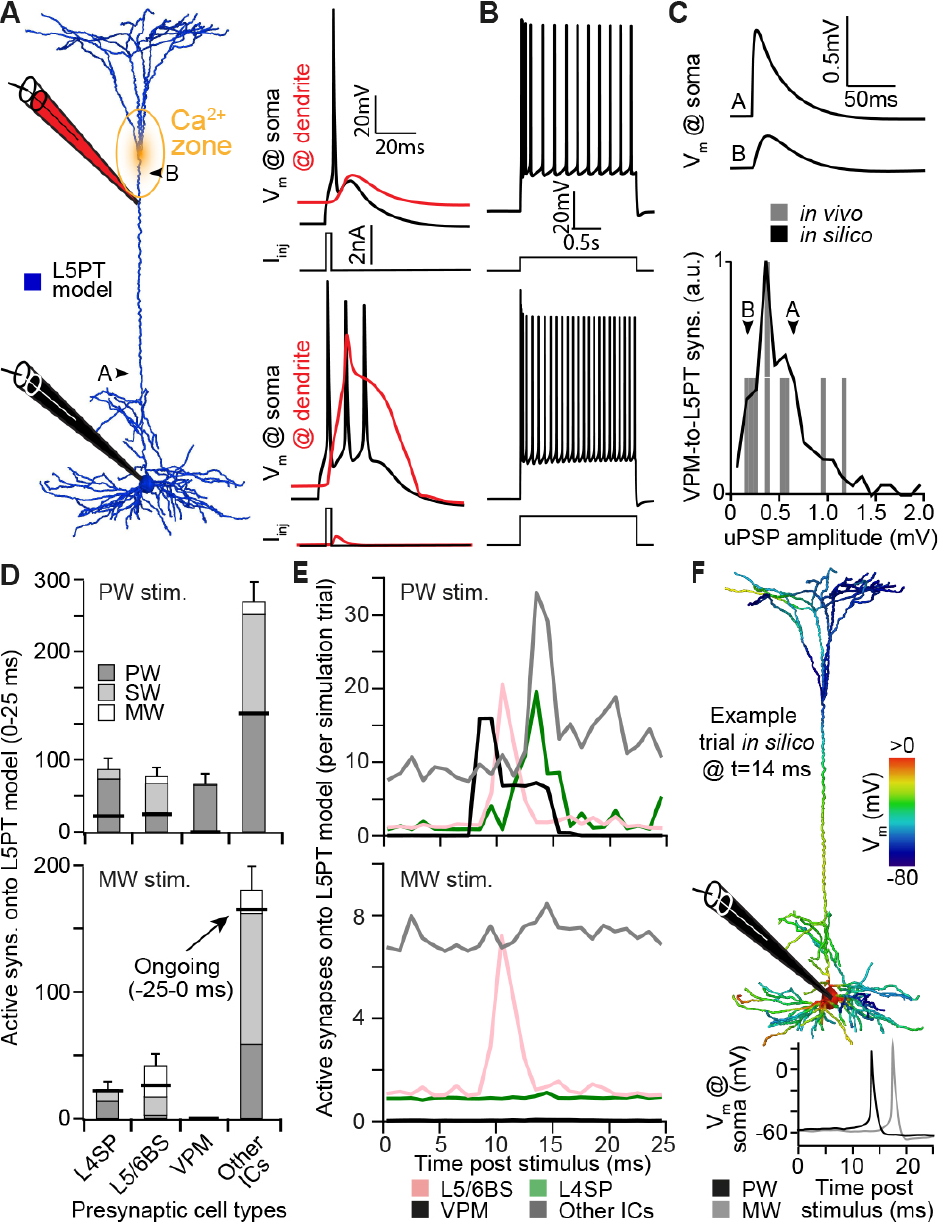
Intrinsic physiological and synaptic constrains for *in silico* experiments. **A)** Left panel: L5PT neuron model, consisting of 1033 dendritic compartments with previously reported biophysical models ^27^. The parameters of the biophysical models were tuned until numerical simulations reproduced current injection-evoked responses that are characteristic for L5PT neurons (right panels): (1) a brief current injection into the soma triggers an AP that back-propagates into the apical dendrite (bAP), and (2) a brief current injection into the soma, followed by current injection into a Ca^2+^ channel dense region around the first bifurcation point of the apical tuft, triggers somatic bursts (i.e., BAC firing). **B)** The model supported regular AP firing of increasing frequencies in response to sustained current injections of increasing amplitude. **C)** The peak conductance at excitatory synapses from different presynaptic cell types was optimized to match empirical unitary post-synaptic-potential (uPSP) amplitude distributions (here exemplified for VPM-to-L5PT synapses ^7^). **D)** The neuron model was embedded into an anatomically realistic network model of rat wS1 and VPM. Based on cell type-specific axo-dendritic overlap (using the morphologies shown in **Fig. 1** and **Fig. S1**) and wRF measurements (**Fig 3** and **Fig. S3**), plausible synaptic input patterns to the L5PT neuron model were generate for deflections of different whiskers. The number (mean ± STD across simulation trials) of active synapses that impinge onto the neuron model during the first 25 ms after simulations of PW and MW deflections are shown. Horizontal black lines denote the number of active synapses that each cell type contributes also to 25 ms of ongoing activity (i.e., before the stimulus). Different grey shadings denote the location of the presynaptic neurons (i.e., somatotopically aligned with the PW, SW or MW). **E)** The number (mean across simulation trials) of active synapses that impinge onto the neuron model during each millisecond of the first 25 ms after simulations of PW and MW deflections, respectively. Other ICs represent inputs from all excitatory intracortical cell types, except for the border stratum cells. **F)** Exemplary simulation trial for synaptic input patterns as shown in panels D and E, which are integrated by the dendrites of the L5PT neuron model (upper panel) and transformed into somatic APs (lower panel).

**Figure S6:**
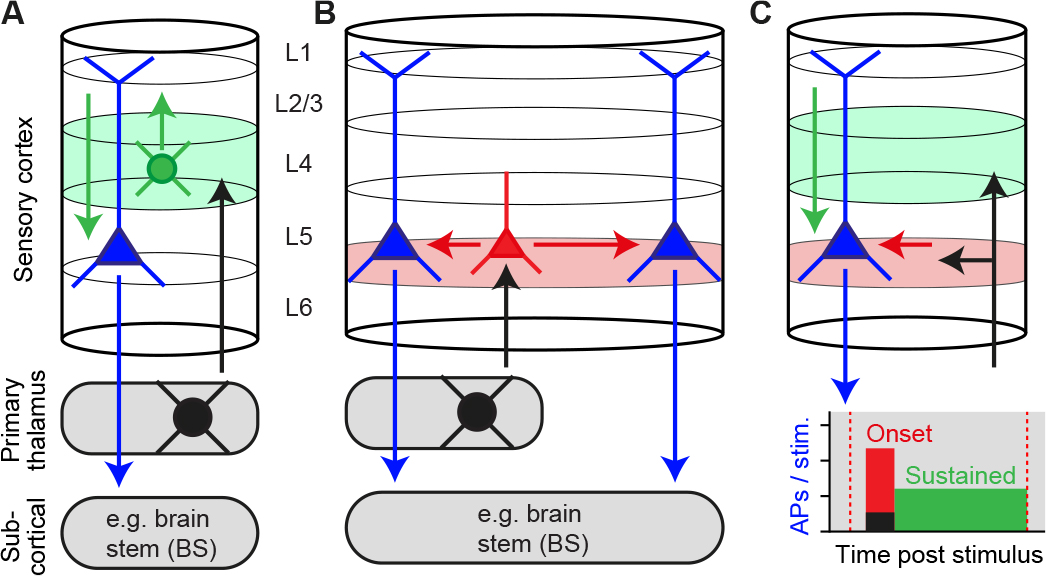
Suggested concept of primary sensory cortex. Sensory-evoked thalamocortical input is relayed in parallel by two orthogonally organized thalamorecipient populations which give rise to complementary canonical pathways: vertical to layers 2/3 by L4SP neurons **(A)**, and horizontally to layers 5/6 by L5/6BS cells **(B)**. The deep thalamorecipient pathway activates pyramidal tract neurons, whereas signal flow in the upper layers terminates in layer 5. The complementary pathway theory hence provides a potential explanation for sustained AP responses in pyramidal tract neurons that persist for the duration of the stimulus. We showed that one way to drive cortical output is by providing sufficiently strong and synchronous synaptic input to the proximal dendrites. However, synchrony decreases during recurrent excitation within local and long-range cortical circuits. Moreover, a substantial fraction of these recurrent and top-down inputs will impinge onto distal dendrites (e.g. within L1). It is hence unlikely that sustained responses in pyramidal tract neurons originate from the same mechanism as the onset responses (see also ^14^). We thus hypothesize that the L5/6BS cell-driven onset responses are required to switch the apical dendrites into an active state, which allows pyramidal tract neurons to transform temporally less synchronous and spatially more distributed synaptic inputs (e.g. from layers 2/3) into sustained patterns.

**Table S1.** Biophysical parameters of the L5PT model. These parameters were obtained using the multi-objective optimization algorithm described previously ^27, 51^. Units for different ion channel densities are pS/μm^2^. τ_Ca_ (ms) is the time constant of the calcium buffering model, and γ_Ca_ is a dimensionless parameter describing the calcium buffer affinity. g_pas_: passive membrane conductance; Na_t_: fast inactivating sodium current; Na_p_: persistent sodium current; K_t_: fast inactivating potassium current; K_p_: slow inactivating potassium current; SKv3.1: fast non-inactivating potassium current; SK E2: calcium-activated potassium current; Ca_LVA_: low voltage-activated calcium current; Ca_HVA_: high voltage-activated calcium current; I_m_: muscarinic potassium current; I_h_: non-specific cation current. * Density in the calcium “hot zone” between 900-1100 μm from the soma. The density of low- and high-voltage activated calcium channels in the apical dendrite was set to 1% and 10% of that value, respectively, outside of the “hot zone”. ** The density of I_h_ in the apical dendrite increases exponentially with distance d to the soma, with parameters A = −0.8696 pS/μm^2^, B = 2.087pS/μm^2^, C=3.6161, and d_max_ the distance of the apical dendrite top located the furthest from the soma. Voltage- and time-dependence of ion channels was modeled using the HH formalism. All corresponding parameters were taken from the literature and have been described in detail previously ^27^.

| Parameter | Soma | AIS / Myelin | Apical dendrite | Basal dendrites |
| --- | --- | --- | --- | --- |
| $C_m$ ($\mu\text{F}/\text{cm}^2$) | 1.0 | 1.0 / 0.04 | 2.0 | 2.0 |
| $r_a$ ($\Omega\text{cm}$) | 100 | 100 / 100 | 100 | 100 |
| $g_{\text{pas}}$ ($1/r_m$) | 0.326 | 0.256 / 0.4 | 0.882 | 0.631 |
| $\text{Na}_t$ | 24300 | 880 / – | 252 | – |
| $\text{Na}_p$ | 49.9 | 14.6 / – | – | – |
| $\text{K}_t$ | 471 | 841 / – | – | – |
| $\text{K}_p$ | 0 | 7730 / – | – | – |
| SKv3.1 | 9830 | 9580 / – | 112 | – |
| SK E2 | 492 | 0.577 / – | 34 | – |
| $\text{Ca}_{\text{LVA}}$ | 46.2 | 85.8 / – | 1040* | – |
| $\text{Ca}_{\text{HVA}}$ | 6.42 | 6.92 / – | 45.2* | – |
| $\tau_{\text{Ca}}$ (ms) | 770 | 507 / – | 133 | – |
| $\gamma_{\text{Ca}}$ (1) | 0.000616 | 0.0175 / – | 0.0005 | – |
| $I_m$ | – | – / – | 1.79 | – |
| $I_h$ | 0.8 | 0.8 / – | $A+B \cdot \exp(C \cdot d/d_{\text{max}})$ ** | 2 |

**Table S2.** Features of membrane potential used to constrain the intrinsic physiology of the L5PT model. Empirical features were adapted as reported previously ^27^. ISI: inter-spike interval; AHP: after-hyperpolarization. Model features based on optimized parameters (see **Table S1**). Difference between model features and average experimental features given in units of STD of the experimental features. The recording locations for the bAP amplitude were adjusted to account for a longer apical trunk of the present L5PT morphology.

| Feature | Mean ± STD | Model | Difference (STD) |
| --- | --- | --- | --- |
| Ca<sup>2+</sup> AP peak | 6.73 ± 2.54mV | 10.8mV | 1.6 |
| Ca<sup>2+</sup> AP width | 37.43 ± 1.27ms | 36.5ms | 0.7 |
| BAC AP count | 3 ± 0 | 3 | 0 |
| Mean somatic AP ISI | 9.9 ± 0.85ms | 9.4ms | 0.6 |
| Somatic AHP depth | -65 ± 4mV | -66mV | 0.3 |
| Somatic AP peak | 25 ± 5mV | 34mV | 1.8 |
| Somatic AP half-width | 2 ± 0.5ms | 1.6ms | 0.8 |
| AP count (somatic current injection only) | 1 ± 0 | 1 | 0 |
| bAP amplitude at 835μm from the soma | 45 ± 10mV | 14mV | 3.1 |
| bAP amplitude at 1015μm from the soma | 36 ± 9.33mV | 9mV | 2.9 |

**Table S3.**
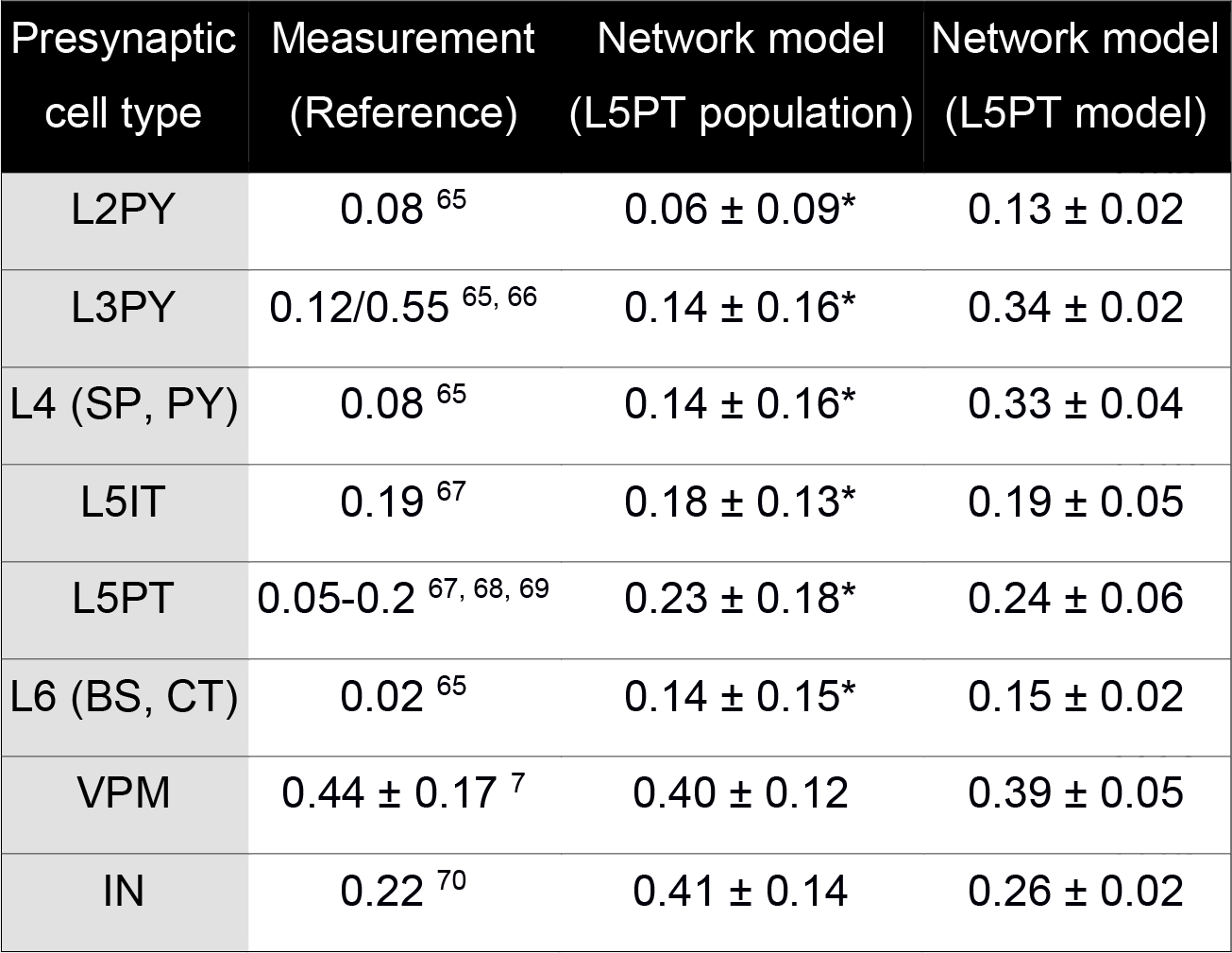
Comparison between predicted connection probabilities in wS1 network model and previously reported measurements from paired-recordings *in vitro* or *in vivo*. The * denotes predicted connection probabilities between truncated morphologies in 300 μm wide thalamocortical/semi-coronal slices of network model.

**Table S4.** Features of uPSP distributions of L5PTs for synaptic input from each presynaptic excitatory cell type, and the respectively fitted synaptic conductance values. Empirical values for uPSP amplitude distributions of synapses from IC cell types ^61^ and VPM thalamus ^7^ were adapted as reported previously. https://www.dropbox.com/s/tg2kl837homq4dp/V11.mp4?dl=0

| Cell type | uPSP Mean (mV)<br>(exp. / fit) | uPSP Median (mV)<br>(exp. / fit) | uPSP Max. (mV)<br>(exp. / fit) | Conductance per<br>synapse (nS) |
| --- | --- | --- | --- | --- |
| L2PY | 0.49 / 0.43 | 0.35 / 0.37 | 1.90 / 2.50 | 1.47 |
| L3PY | 0.49 / 0.44 | 0.35 / 0.39 | 1.90 / 1.98 | 1.68 |
| L4 (SP, PY) | 0.35 / 0.35 | 0.33 / 0.30 | 1.00 / 1.41 | 1.14 |
| L5IT | 0.47 / 0.40 | 0.33 / 0.35 | 1.25 / 1.70 | 1.38 |
| L5PT | 0.46 / 0.43 | 0.36 / 0.39 | 1.50 / 1.46 | 1.59 |
| L6 (BS, CC) | 0.44 / 0.42 | 0.31 / 0.40 | 1.80 / 1.26 | 1.63 |
| L6CT | 0.44 / 0.39 | 0.31 / 0.36 | 1.80 / 1.73 | 1.80 |
| VPM | 0.571 / 0.51 | 0.463 / 0.44 | 1.18 / 1.80 | 1.78 |

**Movie S1:** Examples of *in silico* pharmacology experiments.

